# Repulsive vs. Attractive Crowding Distinctly Regulate TDP-43 Condensates through Region-Specific Structural Dynamics

**DOI:** 10.1101/2025.07.21.665869

**Authors:** Guoqing Zhang, Cibo Feng, Xiakun Chu

**Author notes:** Equal contribution to this work (Guoqing Zhang, Cibo Feng).

## Abstract

TAR DNA-binding protein 43 (TDP-43) is crucial for RNA processing and nucleocytoplasmic transport, and its pathological aggregation is a hallmark of neurodegenerative disorders such as amyotrophic lateral sclerosis and frontotemporal dementia. The intrinsically disordered, prion-like C-terminal domain (CTD) of TDP-43 drives its liquid-liquid phase separation (LLPS), a process fundamental to both normal cellular function and disease-associated aggregation. Using coarse-grained molecular simulations, we systematically explored how distinct macromolecular crowding environments, including repulsive (steric) and attractive (interaction-based) crowding conditions, influence the phase behavior and internal organization of TDP-43 CTD condensates. Our findings reveal that both repulsive and attractive crowders maintain strong correlations between single-chain compaction, dimerization propensity, and macroscopic phase separation, yet operate via distinct physical mechanisms: repulsive crowders drive condensation through volume-exclusion-mediated entropic stabilization, while attractive crowders modulate condensate formation via competitive enthalpic interactions. Spatial structural analysis identifies a robust region-specific internal architecture, with *α*-helices enriched at the condensate core and aligned parallel to the interface, and intrinsically disordered regions (IDRs) preferentially localized near the condensate surface in orientations nearly perpendicular to the interface. Dynamically, *α*-helical regions form strong yet transient interaction hubs, whereas IDRs establish weaker but more persistent contacts, thereby maintaining structural fluidity. Collectively, our results uncover a regulatory principle by which macromolecular crowding modulates TDP-43 condensation through distinct entropic and enthalpic contributions, offering key mechanistic insights into condensate formation and dysregulation relevant to neurodegenerative diseases.

## Introduction

TAR DNA-binding protein 43 (TDP-43) is a multifunctional RNA-binding protein involved in numerous aspects of RNA metabolism, including pre-mRNA splicing, RNA transport, translation regulation, and mRNA stability.^1–3^ Under normal conditions, TDP-43 is predominantly localized in the nucleus. However, its mislocalization to the cytoplasm, followed by aberrant accumulation and aggregation, is a hallmark of several neurodegenerative diseases, most notably amyotrophic lateral sclerosis (ALS) and frontotemporal dementia (FTD).^4–7^ Emerging evidence suggests that pathological TDP-43 aggregation often originates from liquid-liquid phase separation (LLPS),^8,9^ a reversible process through which proteins and nucleic acids condense into dynamic, membraneless organelles. These condensates play essential roles in regulating the spatial and temporal organization of cellular biochemistry.^10^ However, under disease conditions, such phase-separated assemblies can undergo aberrant maturation into irreversible, insoluble aggregates.^11–13^ Despite extensive studies, the molecular determinants that drive the transition from functional LLPS to pathological aggregation remain incompletely understood. ^14^ Elucidating these mechanisms is crucial for understanding the pathogenesis of TDP-43-related disorders and may inform the development of therapeutic strategies aimed at preventing or reversing pathological protein aggregation.^15,16^

Structurally, TDP-43 comprises several distinct functional domains: an N-terminal domain (NTD), two RNA recognition motifs (RRM1 and RRM2), and a C-terminal intrinsically disordered domain (CTD, residues 267-414). ^17^ The CTD is a low-complexity, prion-like region enriched in glycine, glutamine/asparagine, and aromatic residues, and it harbors the majority of ALS-associated mutations. ^18,19^ This disordered region is central to TDP-43’s self-assembly propensity, playing a critical role in both physiological phase separation and pathological aggregation.^8,20–22^ Recent studies have revealed that LLPS driven by the TDP-43 CTD results from a cooperative interplay between distinct interaction modalities: stable, site-specific interactions mediated by a highly conserved *α*-helical segment (residues 319-341), and transient, dynamic interactions involving aromatic residues (e.g., phenylalanine) and aliphatic residues (e.g., methionine) within two flanking intrinsically disordered regions (IDRs), including IDR1 (residues 267-318) and IDR2 (residues 342-414). ^23–25^ This synergy between structural and disordered motifs underlies the dual nature of TDP-43’s condensate behavior, enabling both its functional and pathological assemblies.

It is important to note that most of the mechanistic insights for LLPS have been derived from *in vitro* studies using simplified systems composed of purified proteins and, in some cases, supplemented with synthetic polymers to mimic aspects of the crowded intracellular environment.^26^ However, the cytoplasm of living cells presents a far more complex and heterogeneous milieu, densely populated with macromolecules including proteins, nucleic acids, polysaccharides, and metabolites.^27,28^ This phenomenon, referred to as macromolecular crowding, significantly alters the thermodynamics and kinetics of biomolecular interactions, and has been shown to influence both the formation and the physical properties of biomolecular condensates.^29–33^ Despite significant progress in identifying residue-specific contributions to TDP-43 phase behavior, ^20,23,34,35^ the influence of cellular environmental factors, such as macromolecular crowding, ionic strength, post-translational modifications, and interactions with other biomolecules, on TDP-43 LLPS is not fully understood. ^36,37^ A comprehensive understanding of these environmental modulators is essential to delineate the mechanisms driving both physiological and pathological LLPS,^38^ offering valuable insights into potential therapeutic targets and strategies for TDP-43-associated diseases.

In general, macromolecular crowding is widely recognized to promote LLPS through excluded volume effects, which effectively increase the local concentration of phase-separating proteins and thereby stabilize condensate formation.^39–43^ This entropic mechanism arises from steric repulsion imposed by surrounding macromolecules, which limits the available volume and favors demixing into dense and dilute phases. To emulate the crowded intracellular environment, simplified *in vitro* systems often employ synthetic crowding agents such as polyethylene glycol (PEG), Ficoll, or dextran.^26^ These agents primarily recapitulate excluded volume effects and serve as tractable proxies for the high macromolecular content of the cytoplasm. By modulating their size and concentration, such crowding agents help bridge minimal reconstitution assays with the complex, heterogeneous conditions of living cells.^44,45^

However, the intracellular environment is far from an inert background. Instead, it is a complex, dynamic mixture of proteins, RNAs, and small molecules that can directly or indirectly interact with LLPS-prone proteins. ^46–48^ These cellular constituents may engage in specific or nonspecific interactions with client proteins, potentially competing with or disrupting the transient contacts essential for condensate stability.^11,49–52^ As a result, the net effect of crowding on LLPS depends critically on the physicochemical nature of the crowders and their interaction profiles. Broadly, crowding effects can be simplified as either repulsive (steric) or attractive (interaction-based). In repulsive crowding, crowders are inert to the client protein and promote LLPS through depletion interactions, effectively increasing inter-protein association via excluded volume.^39–43,53^ In contrast, attractive crowders weakly bind or transiently interact with the protein, potentially competing with self-association and altering the energy landscape of phase separation.^54^ These two regimes yield qualitatively distinct outcomes: repulsive crowders enhance LLPS by stabilizing intermolecular contacts and facilitating condensate nucleation, whereas attractive crowders may sequester binding sites, introduce enthalpic penalties, or engage in competing interactions that suppress or rewire phase behavior.^55,56^ In cellular contexts, both types of interactions coexist and jointly determine condensate formation, dynamics, and material properties. For example, while a high total macromolecular content tends to support LLPS, specific interactions with RNAs, chaperones, or co-aggregating proteins can modulate condensate stability or lead to functional or pathological remodeling. ^26,55^ Disentangling these opposing contributions and quantifying their relative roles remains a central challenge in understanding phase behavior *in vivo*.

In this study, we systematically investigated the LLPS behavior of the TDP-43 CTD under physiologically relevant macromolecular crowding conditions using simplified coarsegrained (CG) molecular dynamics (MD) simulations. By comparing the phase separation dynamics of monomeric, dimeric, and multi-chain TDP-43 CTD systems, we identified both shared and distinct features in their responses to crowded environments. Through detailed contact map analyses, we characterized how the concentration and nature of crowding agents, including repulsive and attractive crowders, modulate the phase behavior of TDP-43 CTD. To gain mechanistic insight into region-specific contributions, we partitioned the TDP-43 CTD into three structurally and functionally defined regions: IDR1, the central *α*-helical segment (Helix), and IDR2. We then examined the thermodynamic and kinetic roles of each region in phase separation across different crowding regimes. Our analysis revealed that inter-segment interaction patterns and segmental flexibility collectively shape the condensation process, highlighting distinct structural roles and interaction propensities within the TDP-43 CTD. Together, these findings fundamentally advance our understanding of LLPS under crowded, biologically relevant conditions. Our study elucidates how region-specific features of TDP-43 CTD, in conjunction with external crowding interactions, cooperatively govern phase behavior. Beyond TDP-43, the framework obtained from our work offers a generalizable approach for probing how macromolecular crowding influences the phase behavior of other intrinsically disordered proteins (IDPs) and biomolecular condensates.

## Materials and Methods

### CG MD simulations

#### Hydropathy scale (HPS) Model

We employed an improved version of the HPS CG model, known as the HPS-Urry model, to describe the residue-level interactions within the TDP-43 CTD. ^57–59^ In this model, each residue is represented by a single CG bead, sequentially connected by harmonic springs to form a polymer chain. The non-bonded interactions among beads capture key physicochemical properties of residues, including hydrophobicity, steric repulsion, and electrostatic interactions screened by ionic strength. The total interaction energy governing the protein is given by:^57–59^

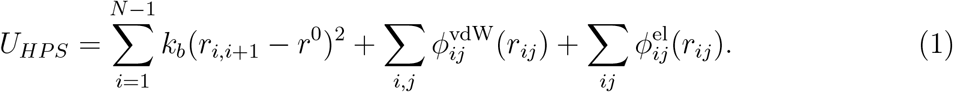

The first term in Eq. 1 represents the harmonic bond potential between adjacent residues (*i, i* + 1) separated by the distance *r*_*i,i*+1_, where *k*_*b*_ = 2000 kcal*/*nm^2^ is the bond spring constant, and *r*^0^ = 0.382 nm is the equilibrium bond length.

The second term in Eq. 1 captures residue-specific van der Waals (vdW) interactions between non-bonded residue pair (*i, j*) separated by distance *r*_*ij*_ and modulated by a hydropathy scale. The interaction is defined as:

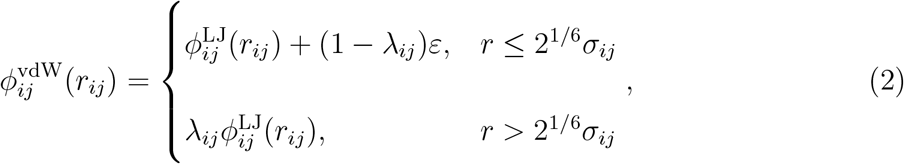

where 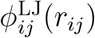 denotes the standard Lennard-Jones (LJ) potential:

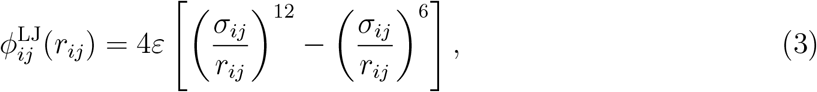

with *ε* = 0.2 kcal/mol representing the interaction energy scale. The parameter *σ*_*ij*_ = (*σ*_*i*_ + *σ*_*j*_)*/*2 defines the effective interaction range, where *σ*_*i*_ is the residue-specific vdW radius of residue *i*. The interaction strength *λ*_*ij*_ between residues *i* and *j* is governed by their relative hydrophobicity and modulates the attractive portion of the potential. These hydropathy parameters were originally derived from partial charges in classical all-atom force fields^60^ and subsequently improved in the HPS-Urry model to better match experimental data.^59^ Specifically, the refined parameter is computed as 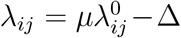, with *µ* = 1.0 and Δ = 0.08, where 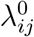 denotes the original hydropathy-derived parameter. The offset Δ was introduced to improve agreement between simulated and experimentally measured *R*_*g*_ for IDPs.^59^ The detailed values of *σ*_*i*_, *λ*_*ij*_ and residue-specific mass can be found in previous studies.^57,59^

The third term in Eq. 1 describes salt-screened electrostatic interactions between charged residues, enabling sequence-specific modeling of electrostatics under physiologically relevant conditions:^61^

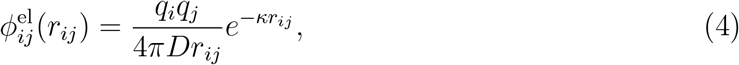

where *q*_*i*_ denotes the integer charges of residues *i*, respectively. Residue charges were assigned based on standard ionization states at physiological pH (pH 7): *q*_*i*_ = +*e* for ARG and LYS, *q*_*i*_ = −*e* for Asp and Glu, and *q*_*i*_ = 0 for HIS and all other residues, assuming neutral protonation states. The dielectric constant of the solvent was set to *D* = 80, approximating that of water, and the inverse Debye screening length *κ* = 1.0 nm^−1^ corresponds to a salt concentration of approximately 100 mM, reflecting typical physiological ionic strength. This Debye-Hückel potential effectively accounts for the exponentially decaying electrostatic potential due to ionic screening in aqueous environments.

#### Crowding interactions

We modeled crowding agents as CG beads, each assigned a mass of 1500 amu and a radius of *r*_*c*_ = 0.8 nm, mimicking the mass and effective size of PEG1500 analogs. Interactions between proteins and crowders 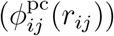 and between crowders themselves 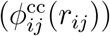 were described using previously established potentials.^48,62^ The total energy contribution from crowding is expressed as:

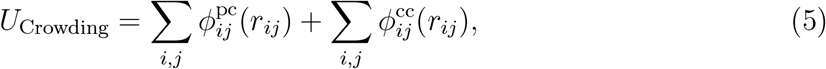

where *r*_*ij*_ denotes the distance between any two CG beads, including both protein residues and crowders.

The protein-crowder interactions are governed by either purely repulsive or LJ-like attractive potentials, depending on the chemical nature of the crowders (repulsive vs. attractive):

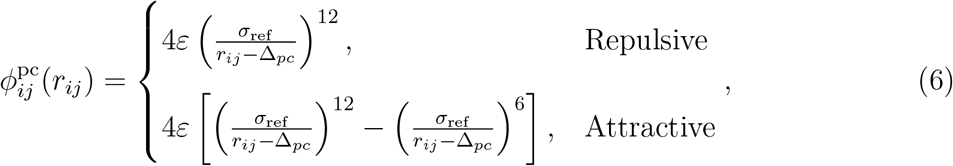

where *σ*_ref_ = 0.6 nm is the reference diameter, and Δ_*pc*_ = *r*^0^*/*2+*r*_*c*_ −*σ*_ref_ = 0.391 nm accounts for the shift between the CG protein bead and the crowder.

Crowder-crowder interactions are modeled purely repulsive, described by:

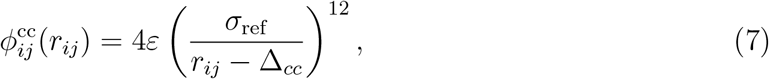

where Δ_*cc*_ = 2*r*_*c*_ − *σ*_ref_ = 1.0 nm.

The volume fraction of crowders (either repulsive *C*_*Rep*_ or attractive *C*_*Att*_), is computed as:

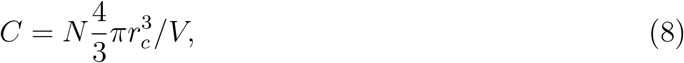

where *N* is the total number of crowder beads and *V* is the volume of the simulation box.

#### Slab simulations for phase coexistence

To efficiently obtain well-converged phase coexistence properties of TDP-43 CTD condensates, we adopted the slab model simulation approach as previously established.^57^ This method accelerates the equilibration of two-phase coexistence by initializing the system in a slab-like configuration, where polymer chains are densely packed within a confined region. Compared to simulations that begin from a homogeneous or randomly dispersed configuration, the slab setup achieves equilibrium significantly faster by providing an initial dense phase.^57^

The simulation procedure involved the following steps. First, 100 TDP-43 CTD chains were randomly placed inside a cubic simulation box. A short NPT run was then performed under an elevated pressure of 100 atm applied in all three directions, allowing the system to compress isotropically. The NPT simulation was terminated once the box reached dimensions of approximately 15 nm *×* 15 nm *×* 15 nm, yielding a compact, slab-like dense phase. The box was subsequently elongated in the *z*-direction to a final size of 15 nm *×* 15 nm *×* 50 nm, and a 5 *µ*s NVT simulation was performed to allow the system to evolve toward a steady-state phase-separated configuration.

To investigate the effect of macromolecular crowding, we introduced CG crowders into the system at defined volume fractions. Repulsive (*C*_Rep_) and attractive (*C*_Att_) crowders were added respectively at concentrations of 5% (N = 262), 10% (N = 525), 15% (N = 787), 20% (N = 1049), 25% (N = 1311), and 30% (N = 1574). Following an additional equilibration phase, we analyzed the trajectories after discarding the initial 1 *µ*s of each simulation, ensuring that only the equilibrated phase behavior was used for subsequent structural and thermodynamic characterization.

#### Simulation protocols

To ensure reliable sampling of the conformational space for both single-chain and dimer systems of TDP-43 CTD, we employed replica exchange molecular dynamics (REMD) simulations.^63^ For single-chain simulations, repulsive and attractive crowders were added at volume fractions of 5% (N = 364), 10% (N = 728), 15% (N = 1092), 20% (N = 1457), 25% (N = 1821), and 30% (N = 2185) within a cubic simulation box of size 25 nm*×*25 nm*×*25 nm. For dimer simulations, the same crowder volume fractions were used, corresponding to 5% (N = 190), 10% (N = 381), 15% (N = 572), 20% (N = 762), 25% (N = 953), and 30% (N = 1144), placed inside a spherical confinement of radius 12.5 nm. The spherical restraint was implemented using PLUMED.^64^

Each REMD simulation was performed over 32 temperature replicas, ranging from 150 K to 800 K. Each replica was simulated for a total of 10 *µ*s, with replica exchange attempted every 1 ns. The average exchange acceptance rate was greater than 31.4% for single-chain simulations and greater than 24.1% for dimer simulations. For analysis, only the final 9 *µ*s of the 300 K replica were used, discarding the first 1 *µ*s as equilibration.

All simulations, including REMD and slab-based LLPS simulations, were performed using the LAMMPS package^65,66^, with the real unit system. The *α*-helical regions of TDP-43 CTD were treated as rigid bodies in all simulations. A timestep of 10 fs was used, with output written every 100 ps. Langevin dynamics was applied at 300 K, using a damping constant of 1000 ps for LLPS simulations and 1 ps for single-chain and dimer REMD simulations. Non-bonded interactions were truncated at 3.5 nm, and periodic boundary conditions were applied in all three spatial directions.

### Quantities calculations

#### Radial density function (RDF)

To characterize the spatial distribution of crowders around the TDP-43 CTD, we calculated the RDF by counting the number of crowder beads within spherical shells centered at the protein’s center of mass. The radial distribution function *P* (*r*) at a distance *r* is given by:

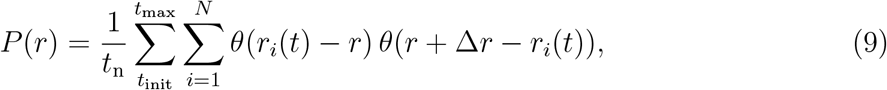

where *t*_max_ is the total simulation time, *t*_init_ time prior to equilibration (1 *µs*) that was discarded from the analysis, *t*_*n*_ = *t*_max_ − *t*_ini_ is the number of analyzed simulation frames, *N* denotes the total number of crowder beads, and *r*_*i*_ is the distance of the *i*th crowder bead from the center of mass of the TDP-43 CTD. The shell thickness was set to Δ*r* = 0.1 nm. The Heaviside step function *θ*(*x*) is defined as:

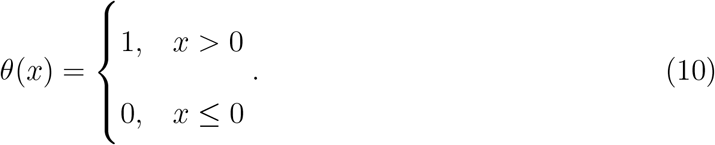

The RDF *ρ*(*r*) was then obtained by normalizing the number distribution with the volume of the shell:

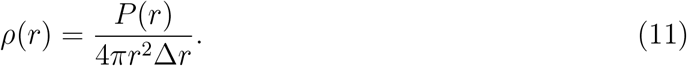

#### Mean square displacement (MSD)

We calculated the MSD of TDP-43 CTD within condensates using the following expression:

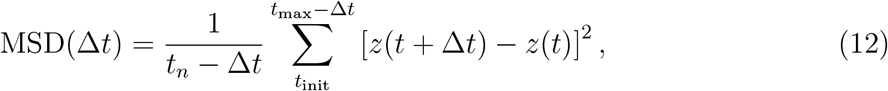

where Δ*t* is the lag time, *z*(*t*) denotes the position of the chain’s center of mass in the *z*direction at time *t*. The MSD was calculated as a time average over the simulation trajectory after equilibration. The MSD was further averaged over 100 TDP-43 CTD chains to account for ensemble variability.

To extract the diffusion coefficient *D*, we fit the MSD to a power-law model:

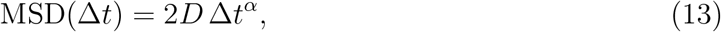

where *α* characterizes the diffusion behavior (*α* = 1 for normal diffusion, *α <* 1 for subdiffusion). To estimate *α*, we performed a log-log fit of the MSD:

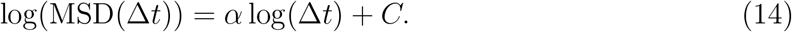

The extracted value of *α* was then inserted back into Eq. 13 to fit the diffusion coefficient *D*.

#### Residue-based contact number

We calculated the contact number (*N*_*c,ij*_) between residues *i* and *j* of TDP-43 CTD using a distance-based cutoff criterion applied over the simulation trajectory after equilibration:

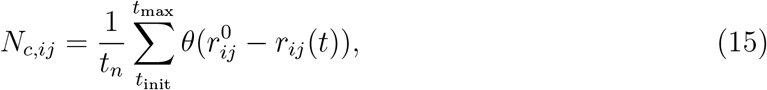

where *r*_*ij*_(*t*) is the instantaneous distance between residues *i* and *j* at time *t*, and the cutoff distance 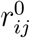 was defined as 0.75(*σ*_*i*_ + *σ*_*j*_), where *σ*_*i*_ and *σ*_*j*_ are the effective vdW radii of residues *i* and *j*, respectively.

For inter-chain contact analysis, we ensured that residues *i* and *j* belonging to different TDP-43 CTD chains. For the phase-coexistence simulations, contact maps were calculated by averaging over 100 chains for intra-chain contacts and over 4950 chain pairs for inter-chain contacts.

### Region-based orientation angle

To characterize the orientation of the three segments (Helix, IDR1, and IDR2) in TDP-43 CTD, we calculated the angular deviation of a segmental vector relative to the z-axis. For each segment of length *L* residues, we defined a representative vector *r* by randomly selecting two residues within the segment:

- For the Helix and IDR1 segments: randomly select residues from the second half 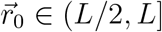 and the first half 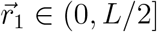, respectively.
- For the IDR2 segment: randomly select residues from the first half 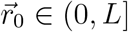 and the second half 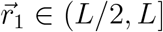, respectively.

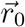 and 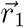 are the 3D coordinates of the selected residues. The orientation vector is then calculated as:

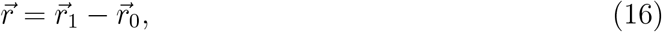

The angle *ϕ*_*k*_ between the segment vector 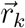 of the *k*-th chain segment and the z-axis unit vector 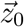 is given by:

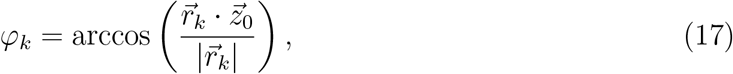

Where 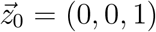.

For each segment, we also calculated the z-coordinate of its center of mass and constructed a 2D histogram of *ϕ* versus z-position.

To define the interfacial region, we identified the range of z-values where the spatial probability density falls between 0.2 and 0.8 times the maximum. Segment orientations within this surface region were extracted and visualized as violin plots, with the left and right halves representing orientations on opposite sides of the condensate surface.

#### Region-based contact number and relaxation time

Based on the residue-level inter-chain contact number 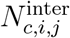 (as defined in Eq. 15), we further calculated the average inter-chain contact number 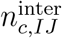 between two different segments *I* and *J*, of lengths *a* and *b*, respectively (where *I /*= *J*). The segments considered are IDR1, IDR2, and Helix:

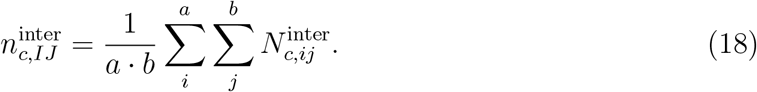

To quantify the dynamics of inter-segment contacts, we calculate the normalized autocorrelation function of 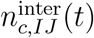 over the simulation trajectory after equilibration:

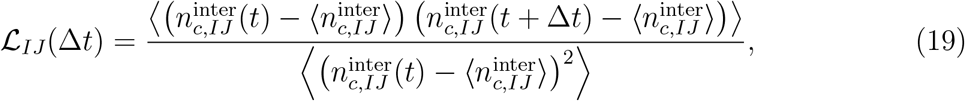

where ⟨ · ⟩ denotes a time average.

To extract the characteristic contact relaxation time *τ*, we fit the autocorrelation function to an exponential decay:

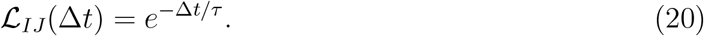

## Results

### TDP-43 CTD phase behaviors under varying crowding conditions

We performed CG MD simulations of the TDP-43 CTD under a range of macromolecular crowding conditions to dissect how purely repulsive versus attractive crowders influence its phase separation behavior. Using the HPS-Urry model to represent the sequence-specific interactions of TDP-43 CTD,^57,59^ we introduced crowding agents as single-bead CG beads at volume fractions ranging from 0% to 30% (repulsive crowder volume fraction denotated as *C*_Rep_ or attractive crowder volume fraction denotated as *C*_Att_). Repulsive crowders were modeled with purely excluded-volume interactions, while attractive crowders interacted nonspecifically with the proteins, as described previously^48^ (see Materials and Methods for details). Strikingly, the presence of crowders significantly altered TDP-43 CTD condensate formation in a manner dependent on crowder type (Figure 1A). In the absence of crowders, TDP-43 CTD chains readily underwent phase separation, consistent with their intrinsic propensity for homotypic association reported in previous studies.^23,25^ Under repulsive crowding conditions, condensate formation was further promoted: nearly all TDP-43 CTD chains coalesced into a single, compact droplet. In contrast, attractive crowders disrupted this self-association; condensates were less compact and more dispersed, with crowders intermixed among the TDP-43 CTD chains. These simulation snapshots illustrate the distinct condensate morphologies promoted by different crowding regimes: repulsive crowders are excluded from the dense phase, enhancing protein-protein interactions, whereas attractive crowders infiltrate the condensate, weakening homotypic contacts and diluting the assembly (Figure 1A).

**Figure 1:**
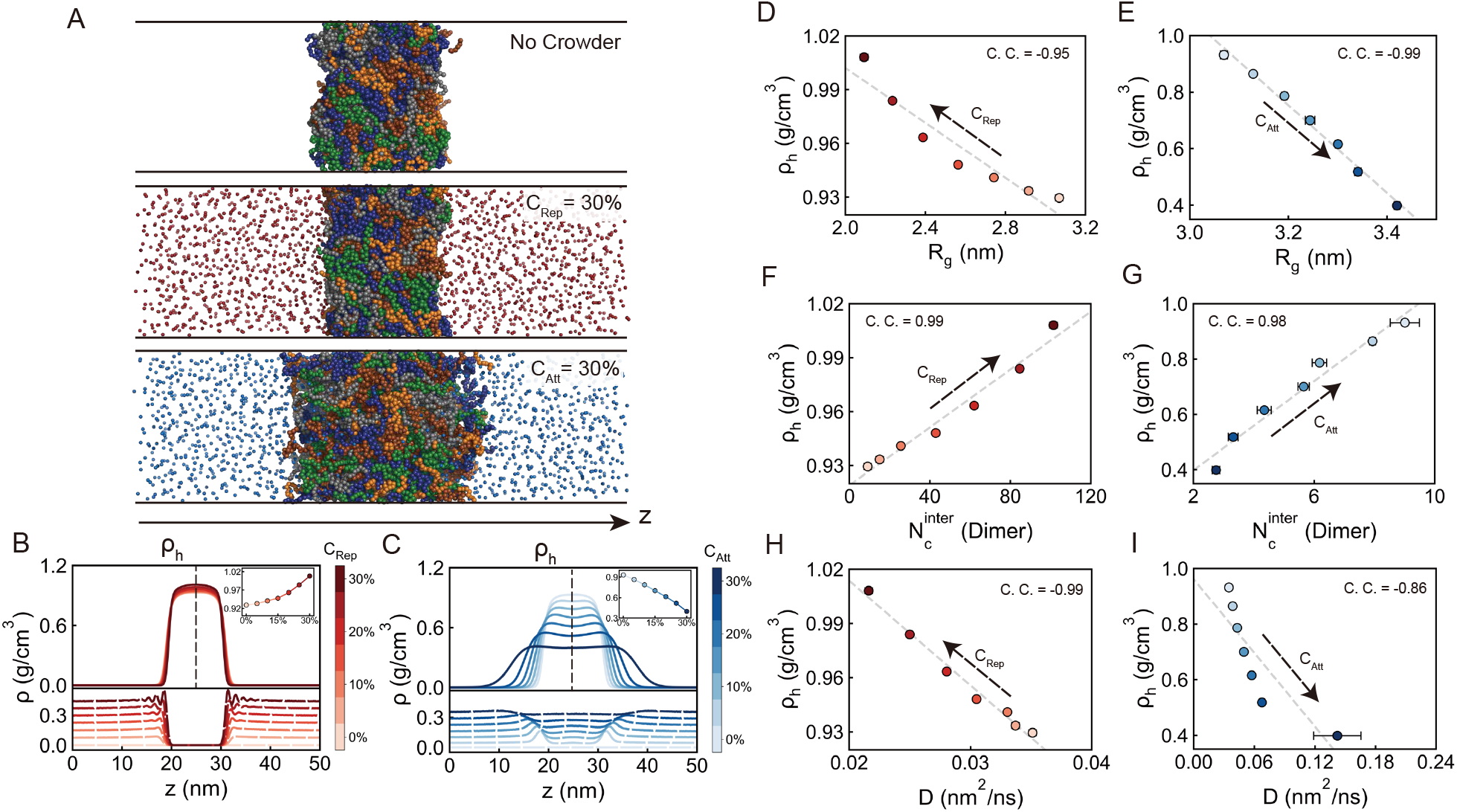
Effects of repulsive and attractive crowders on TDP-43 CTD condensate formation and associated single-chain and dimer-level properties. (A) Representative simulation snapshots showing condensate morphology under three conditions: no crowders, repulsive crowders (30% volume fraction, *C*_Rep_=30%, represented by red beads), and attractive crowders (30% volume fraction, *C*_Att_=30%, represented by blue beads). (B, C) Density profiles along the *z*-axis for TDP-43 CTD (solid lines, upper panels) and crowders (dashed lines, lower panels) in systems with varying concentrations of repulsive (*C*_Rep_) and attractive (*C*_Att_) crowders, respectively. *ρ*_*h*_ denotes the protein density at the condensate core. Insets show the dependence of *ρ*_*h*_ on crowder concentration. (D, E) Relationship between *ρ*_*h*_ and the radius of gyration (*R*_*g*_) of isolated TDP-43 CTD chains across different *C*_Rep_ and *C*_Att_ values. (F, G) Relationship between *ρ*_*h*_ and inter-chain contact number 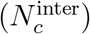 for TDP-43 CTD dimers across different *C*_Rep_ and *C*_Att_ values. (H, I) Relationship between *ρ*_*h*_ and diffusion coefficient (*D*) within the dense phase across different *C*_Rep_ and *C*_Att_ values. In panels (D-I), data points represent means *±* standard errors computed from 5 equal partitions of the total dataset, with Pearson correlation coefficient (C. C.) reported. Dashed lines are provided as visual guides to highlight the trends.

To quantitatively assess how varying crowder concentrations affect TDP-43 CTD condensates, we calculated density profiles for both the proteins and the crowders (Figures 1B and 1C). We observed that increasing concentrations of repulsive crowders systematically elevated the protein density within the condensate core, whereas increasing concentrations of attractive crowders led to a progressive reduction. These trends are consistent with prior computational and experimental studies on crowding-induced modulation of biomolecular condensates.^55,67,68^ Mechanistically, the density enhancement under repulsive crowding arises through two distinct regimes depending on the crowder concentration. At low repulsive crowder fractions (low *C*_Rep_), volume exclusion dominates: sparsely distributed crowders exert modest steric hindrance, but nonetheless drive proteins to partition into the dense phase to minimize the entropic cost of reduced accessible volume. This results in protein depletion from the dilute phase and a corresponding increase in condensate density. At higher *C*_Rep_, strong steric repulsion from densely packed crowders near the condensate boundary compresses the condensate, further increasing the local protein concentration (Figure 1B). In contrast, attractive crowders exhibit a qualitatively different behavior. These crowders preferentially accumulate at the condensate interface and partially infiltrate the dense phase due to favorable protein-crowder interactions. This co-condensation effect dilutes the protein-rich interior, as crowders compete for interaction sites, weakening protein-protein contacts and lowering the dense-phase protein concentration (Figure 1C). Thus, while repulsive crowders enhance condensation via excluded-volume and steric repulsion effects, attractive crowders reduce condensate density through enthalpic mechanisms of competitive binding.

To quantify the intramolecular structural responses of TDP-43 CTD to macromolecular crowding, we calculated the radius of gyration (*R*_*g*_) of individual chains in both condensed and monomeric states (Figure S1). Strikingly, within the condensate phase, *R*_*g*_ remained largely invariant across a broad range of crowder concentrations and interaction types (repulsive vs. attractive). This robustness suggests that the internal conformation extensions of IDPs are highly resilient to external perturbations once embedded within condensates. Consistent with our previous findings,^69^ this observation supports the notion that condensate-resident IDPs retain their characteristic structural features despite variations in environmental conditions such as temperature, ionic strength, or macromolecular crowding. In contrast, monomeric TDP-43 CTD chains exhibited marked sensitivity to crowding, displaying opposite trends under repulsive and attractive conditions (Figure S1). Under repulsive crowding, *R*_*g*_ decreased monotonically with increasing crowder concentration, indicating progressive chain compaction. This behavior aligns with classical excluded-volume theory: crowding reduces the available conformational space, imposing entropic constraints that favor collapsed configurations to minimize free energy.^70,71^ Conversely, attractive crowders elicited a reversed response, where increasing concentrations led to monotonic chain expansion. This trend reflects an enthalpic mechanism in which favorable protein-crowder interactions effectively solvate the protein, stabilizing more extended conformations. Together, these contrasting *R*_*g*_ responses highlight fundamental differences in how repulsive and attractive crowding modulate IDP conformational ensembles. While repulsive crowders promote chain compaction through entropic confinement, attractive crowders introduce enthalpic effects that drive chain expansion. These findings underscore the importance of distinguishing the specific physicochemical nature of crowding interactions when interpreting IDP behavior in both cellular and synthetic environments.

To elucidate the connection between single-chain conformational properties and phase separation propensity under crowding, we examined the relationship between the *R*_*g*_ of isolated TDP-43 CTD chains and the dense-phase protein concentration (*ρ*_*h*_) in the corresponding condensates (Figures 1D and 1E). Under repulsive crowding conditions, increasing crowder concentration induced a progressive decrease in *R*_*g*_, indicative of intramolecular compaction, accompanied by a monotonic increase in *ρ*_*h*_. This coupled behavior suggests that excluded-volume effects simultaneously promote chain collapse and enhance phase separation. These results are consistent with previous studies showing that compact disordered chain conformations are more prone to LLPS, as they facilitate intermolecular association through reduced conformational entropy and increased effective valency.^69,72–75^ Interestingly, in the presence of attractive crowders (Figure 1E), although *R*_*g*_ exhibited an inverse trend of expanding with increasing crowder concentration, the relationship between *R*_*g*_ and *ρ*_*h*_ still remained approximately linear. The strong correlation observed under both repulsive and attractive conditions underscores a key principle: despite the opposing mechanistic origins (entropic compaction vs. enthalpic expansion), single-chain conformational properties such as *R*_*g*_ continue to serve as reliable predictors of phase behavior across diverse crowding environments. This reinforces the notion that molecular-level structural parameters are deeply coupled to condensate-level properties, regardless of whether crowding is primarily driven by volume exclusion or by weak, nonspecific interactions.

To further elucidate the mechanisms underlying the divergent effects of repulsive and attractive crowders, we analyzed the radial distribution function (RDF) of crowders relative to the center of mass of the TDP-43 CTD for single-chain systems (Figure S2; see Materials and Methods for details). Based on the RDF profiles, crowders were classified into two distinct spatial categories: (1) internal crowders, located within the average *R*_*g*_ of the protein, and (2) external crowders, positioned outside this radius yet still near the protein surface. Our analysis revealed a strong positive correlation between the fraction of internal crowders and the *R*_*g*_ of the protein chain, suggesting that the presence of crowders within the protein interior critically influences its conformational response. This finding underscores the profound impact of the spatial positioning of crowders on driving protein compaction or expansion.^76^ Importantly, repulsive and attractive crowders mediate protein structure through fundamentally distinct mechanisms. Under repulsive crowding conditions, crowders are predominantly excluded from the protein interior, exerting primarily entropic effects by constraining the accessible conformational space and thereby promoting protein compaction through volume exclusion. Conversely, attractive crowders can infiltrate the protein core and transiently interact with internal residues. These favorable enthalpic interactions effectively compete with intrinsic intra-chain contacts, disrupt the native contact network, and result in chain expansion.

We next investigated how macromolecular crowding modulates intermolecular interactions between TDP-43 CTD chains by quantifying the average inter-chain contact number 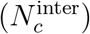 in a dimer system (Figures 1F and 1G). Under repulsive crowding conditions, 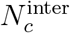 increased significantly compared to the no-crowder baseline, in line with the elevated condensate density (*ρ*_*h*_). This trend indicates that repulsive crowders enhance inter-chain association by promoting close spatial confinement via excluded-volume effects. Such entropic compression increases the likelihood of protein-protein encounters, thereby stabilizing homotypic contacts. In contrast, attractive crowders markedly reduced 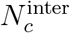, suggesting that the effective stability of protein-protein interfaces is diminished in these environments. This reduction likely arises because attractive crowders compete with TDP-43 CTD for binding sites, disrupting native inter-chain interactions and favoring protein-crowder contacts instead. To further probe how intermolecular contact formation translates to macroscopic phase behavior, we examined the correlation between 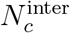 and *ρ*_*h*_ across varying crowder concentrations. Under both repulsive and attractive conditions, this correlation remained strong and monotonic, reinforcing the established link between enhanced dimer interactions and increased phase separation propensity.^73,77^ However, these findings underscore a key mechanistic distinction: repulsive crowders promote LLPS by strengthening homotypic interactions via steric confinement and entropic depletion, whereas attractive crowders can attenuate LLPS by redirecting contacts toward heterotypic, often destabilizing, interactions. This highlights the necessity for predictive models of LLPS to incorporate not only crowder concentration but also the physicochemical nature and interaction specificity of crowding agents.^73,77^

We further analyzed the chain dynamic properties of TDP-43 CTD condensates by quantifying the diffusion coefficient (*D*) of proteins under varying crowding conditions (Figures 1H and 1I and S3; see Materials and Methods for details). In repulsive crowding environments, *D* decreased monotonically with increasing crowder concentration, indicating progressively hindered molecular mobility. This diffusion slowdown is attributed to enhanced molecular packing and increased viscosity within the dense phase, as repulsive crowders compress the condensate and restrict internal motions. In contrast, attractive crowders exerted a different influence: the diffusion coefficient increased with crowder concentration. This acceleration is likely due to reduced condensate density, as strong protein-crowder interactions outcompete protein-protein interactions and lead to looser molecular packing. Under these conditions, individual TDP43-CTD molecules experience less confinement and greater translational freedom within the dense phases. Thus, repulsive and attractive crowders modulate protein diffusion within condensates via fundamentally distinct mechanisms. Repulsive crowders enhance physical confinement through excluded volume effects, reducing mobility, whereas attractive crowders increase mobility by dynamically sequestering proteins and disrupting homotypic interactions (Figures 1H and 1I). This duality is consistent with crowding theory, wherein diffusion scales inversely with effective volume occupancy and is modulated by interaction-mediated friction. ^78^ Overall, our findings underscore the importance of protein-crowder interactions in regulating not only condensate architecture but also the internal molecular dynamics of biomolecular condensates.

In summary, our study demonstrates that repulsive and attractive crowding agents exert fundamentally distinct regulatory effects on the phase behavior and internal organization of TDP-43 CTD condensates. Repulsive crowders, mimicking inert volume-excluding macromolecules, promote the formation of compact, tightly networked condensates by enhancing intramolecular compaction and stabilizing inter-chain interactions. These condensates exhibit high density and reduced molecular mobility. In contrast, attractive crowders, which emulate weakly interacting cosolutes, disrupt homotypic contacts and preserve extended chain conformations, resulting in dilute, less cohesive condensates with greater internal dynamics. These opposing behaviors highlight how crowding agents can modulate the energetic and kinetic landscape of phase separation through both entropic and enthalpic contributions.

### Differentiating contact maps of TDP-43 CTD under varying crowding conditions

To elucidate the structural reorganization of TDP-43 CTD under varying crowding conditions, we calculated both intra- and inter-chain residue-level contact probability maps in the absence of crowders, under repulsive crowding, and under attractive crowding (Figure 2, Figures S4-S7; see Materials and Methods for details). These contact maps allow us to dissect how excluded-volume effects and crowder-induced interactions reshape the molecular interaction landscape of TDP-43 CTD across different environments. Under crowder-free conditions, the intra-chain contact map serves as a baseline, revealing a contact hierarchy dominated by local interactions between residues within individual IDRs, with only limited long-range contacts between these two IDRs (Figure 2A). In other words, the two IDRs flanking the central *α*-helical region display extensive intra-region interactions, whereas inter-IDR interactions are sparse. This modular interaction pattern is consistent with previous observations and reflects the intrinsic architecture of the TDP-43 CTD.^12^ Upon phase separation in the absence of crowders, intra-chain contact patterns remain largely conserved, albeit with a modest reduction in the total number of contacts. This reduction suggests a slight conformational expansion of TDP-43 CTD chains within the condensed phase, consistent with increased conformational heterogeneity often observed in phase-separated IDPs.^23^

**Figure 2:**
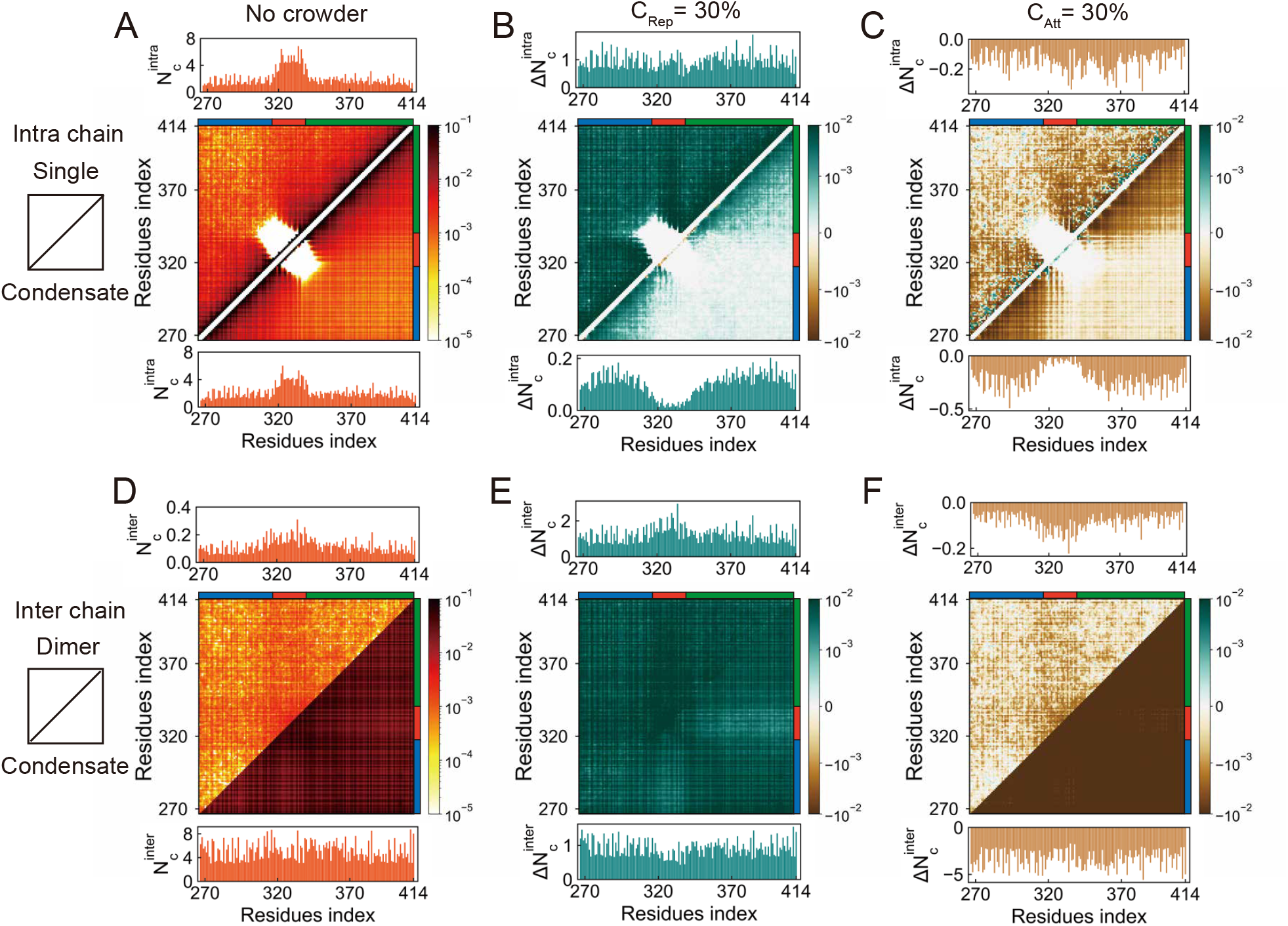
Residue-residue contact probability maps of TDP-43 CTD under different crowding conditions. (A) Intra-chain contact maps for isolated single TDP-43 CTD chains and for individual chains within condensates under crowder-free conditions. (B-C) Differential intra-chain contact maps showing changes in contact probabilities relative to the crowder-free case: (B) with 30% volume fraction of repulsive crowders and (C) with 30% volume fraction attractive crowders. Each panel includes both single-chain and condensate configurations. (D-F) Inter-chain contact probability maps for TDP-43 CTD dimers and for chains within condensates, organized analogously to (A-C). In each contact map, residue indices are shown on both axes. The upper triangle represents the reference configuration (e.g., single chain or dimer), while the lower triangle shows the corresponding condensate configuration. Per-residue contact probabilities (summed across sequence) are displayed above (for the reference) and below (for the condensate) each contact matrix to highlight regions enriched in interactions. The three regions of the TDP-43 CTD, i.e., IDR1, Helix, and IDR2, are indicated as stripes near the axes of each contact map and are colored in blue, red, and green, respectively.

Under repulsive crowding, the intra-chain contact probabilities of single TDP-43 CTD chains increase significantly across the entire contact map, indicating global chain compaction driven by excluded-volume effects (Figures 2B and S4). Sequence-distant regions that seldom interact under dilute conditions are brought into closer proximity due to the reduced accessible volume, resulting in the emergence of long-range contacts. This compaction is quantitatively supported by the observed reduction in the *R*_*g*_ of the TDP-43 CTD (Figure S1). In contrast, TDP-43 CTD chains within the phase-separated condensates exhibit enhanced contact probabilities primarily within individual IDR segments, with little increase in interactions between the flanking IDRs. Thus, even under repulsive crowding, the interaction hierarchy remains preserved: intra-IDR contacts dominate, while inter-IDR contacts remain relatively weak. This structural organization is robust across a wide range of crowder concentrations (Figure S4), suggesting that repulsive crowders uniformly amplify pre-existing interaction motifs rather than triggering novel structural rearrangements. Collectively, these findings indicate that repulsive crowding promotes intramolecular compaction via nonspecific spatial confinement, while largely maintaining the intrinsic contact topology of the TDP-43 CTD. The monotonic, concentration-dependent enhancement of contact strengths across all major intra-chain interaction regions underscores the thermodynamic resilience of TDP-43 CTD’s internal organization in the face of steric crowding.

By contrast, attractive crowders elicit context-dependent structural responses that are essentially opposite to those observed under repulsive conditions, as revealed by our contact map analysis (Figures 2C and S5). In the single-chain regime, we observed an overall reduction in contact probabilities, particularly within the IDRs, indicating that attractive crowders preferentially destabilize intra-IDR contacts. This disruption, especially of inter-IDR interactions, contributes to a notable increase in the *R*_*g*_ (Figure S1). However, within condensates, attractive crowders do not significantly enhance inter-IDR interactions, and the overall pattern of long-range intra-chain contacts remains largely unchanged. This limited restructuring is consistent with the nearly invariant *R*_*g*_ of TDP-43 CTD chains within condensates under attractive crowding conditions (Figure S1). These observations highlight a key mechanistic distinction: while attractive crowders can remodel the conformational landscape of isolated chains by disrupting native intra-chain contacts and promoting chain expansion, their impact within condensates is markedly attenuated, likely due to the prevailing dominance of protein-protein interactions over protein-crowder associations in the densely packed phase.

By examining the inter-chain contacts formed between two TDP-43 CTD chains, our analysis reveals significant differences in inter-chain interaction patterns between dimer and phase-separated states, even in the absence of crowders (Figure 2D). The condensed environment of phase separation enhances inter-chain contacts by approximately an order of magnitude relative to dimer configurations. Both systems share a common hierarchy of interaction strengths: Helix-Helix contacts dominate, followed by Helix-IDR and IDR-IDR interactions. In details, the dimer systems exhibit stronger Helix-IDR than IDR-IDR contacts, while the condensed phase reverses this relationship, with multivalent IDR-IDR interactions becoming dominant. This shift underscores the increasing importance of cooperative, multivalent IDR interactions in stabilizing high-density, phase-separated assemblies.

The introduction of repulsive crowders preserves the fundamental hierarchy of inter-chain interactions across both dimeric and phase-separated systems (Figures 2E and S6). In both configurations, Helix-Helix contacts remain dominant, with condensates exhibiting saturated interactions due to the elevated local concentration of structured helices. Interestingly, dimers under repulsive conditions maintain stronger Helix-IDR contacts than their condensate counterparts, reminiscent of the “fly-casting” mechanism,^61,79^ where flexible IDRs facilitate transient, exploratory interactions with structured domains. This contrast highlights the influence of organizational context on interaction preferences: whereas condensates stabilize Helix-Helix interactions through multivalent avidity, isolated dimers exploit IDR flexibility to explore alternative binding configurations. The persistence of these interaction patterns under repulsive crowding underscores the robustness of intrinsic interaction thermodynamics to nonspecific, excluded-volume perturbations. In contrast, attractive crowders induce opposite changes in inter-chain interactions. In both dimeric and phase-separated systems, increasing crowder concentration consistently reduces inter-chain contact probabilities, correlating with diminished condensate density and weakened homotypic interactions (Figures 2F and S7). Attractive crowders effectively compete with direct residue-residue contacts by favoring crowder-protein interactions, thereby disrupting native inter-chain binding networks.

Our comprehensive contact map analysis reveals fundamentally distinct modes of interaction modulation induced by repulsive versus attractive macromolecular crowding. Under both crowding conditions, single chains, dimers, and phase-separated condensates exhibit monotonic yet opposing responses: repulsive crowders consistently enhance intra- and inter-chain contacts, whereas attractive crowders systematically weaken these interactions. This uniform trend preserves intrinsic interaction hierarchies, allowing reliable prediction of condensate properties such as core density directly from the conformational characteristics of individual molecules (Figures 1D-1I). These observations highlight a crucial mechanistic distinction: repulsive crowders primarily act through uniform volume-exclusion effects, whereas attractive crowders introduce heterogeneous, concentration-dependent interactions that can significantly reorganize the protein contact network. Such complex modulation underscores the delicate interplay between homotypic protein-protein interactions and heterotypic protein-crowder associations, which collectively determine condensate stability and morphology in physiologically crowded environments. ^80^ Ultimately, these findings illuminate the dual functionality of crowders as either stabilizers or antagonists of protein assembly, emphasizing the nuanced balance that governs biomolecular condensate regulation in cellular contexts.

### Region-specific spatial and orientational distributions of TDP-43 CTD within condensates

To elucidate the internal organization of TDP-43 CTD within condensates, we analyzed the spatial distributions and orientation angles of its three structurally defined segments, including the central *α*-helical region and two IDRs (IDR1 and IDR2), along the *z*-axis under different crowding conditions (Figure 3A). Specifically, we compared condensates formed in crowder-free environments, in the presence of purely repulsive crowders, and in the presence of attractive crowders (Figures 3, S8, and S9). This comparative analysis allows us to distinguish how structured versus disordered segments localize and orient within condensates in response to distinct physical perturbations. By disentangling entropic effects arising from excluded-volume interactions from enthalpic effects associated with direct protein-crowder binding, our framework provides mechanistic insights into how diverse crowding conditions reshape the internal architecture and interfacial organization of biomolecular condensates.

**Figure 3:**
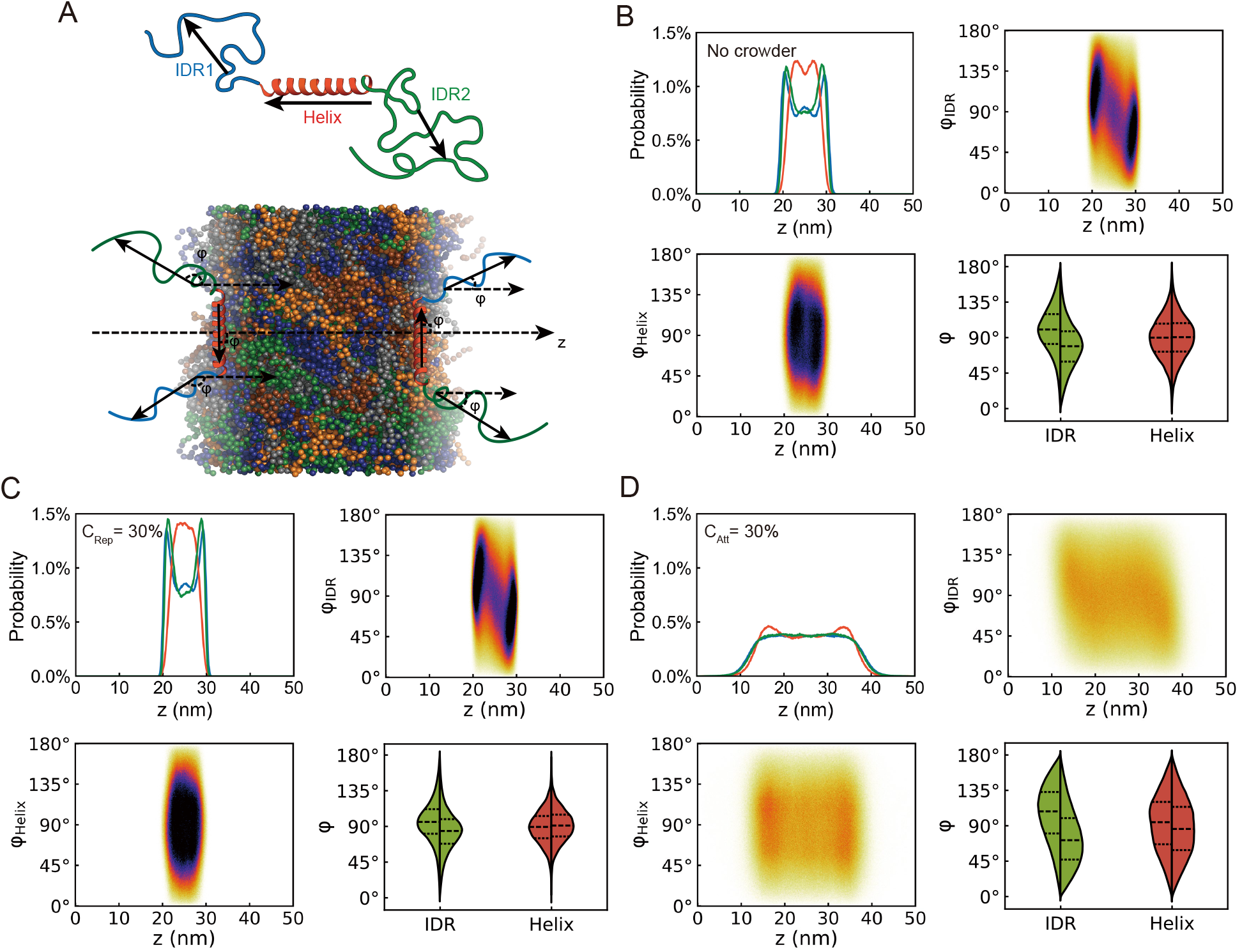
Spatial localization and orientational behavior of TDP-43 CTD segments within condensates under different crowding conditions. (A) Schematic representation of the three regions of TDP-43 CTD: IDR1 (blue), central *α*-helical segment (orange), and IDR2 (green). For each segment, an orientation vector is introduced to calculate its alignment angle relative to the *z*-axis (see Materials and Methods for details). (B-D) Region-specific spatial probability distributions along the *z*-axis and orientation angle distributions of TDP-43 CTD segments within condensates formed under (B) crowder-free conditions, (C) 30% volume fraction of repulsive crowder conditions, and (D) 30% volume fraction of attractive crowder conditions. In each panel of (B-D), the bottom-right subfigure displays the angular distribution for the *α*-helix and IDRs at the left and right surfaces of the condensate, respectively.

In the absence of crowders, TDP-43 CTD chains adopt a stratified internal architecture within condensates (Figure 3B). The central *α*-helical region preferentially localizes toward the condensate core, consistent with its role in mediating specific Helix-Helix interactions that stabilize the dense phase. ^25,81^ In contrast, the flanking IDR1 and IDR2 segments are more broadly distributed along the *z*-axis and tend to localize closer to the condensate periphery. This spatial arrangement suggests a degree of region-level segregation: the structured helix is embedded in the interior to maximize stabilizing interactions, while the flexible IDRs remain more solvent-exposed, dynamically sampling the interfacial region. Angular distribution analysis reveals distinct orientation preferences among the three segments relative to the condensate surface. The *α*-helix exhibits a pronounced tendency to align parallel to the interface, with a peak near 90^*°*^ relative to the *z*-axis. In contrast, the IDRs display mirror-symmetric, surface-specific orientation patterns: on the left surface, both IDRs show preferred angles greater than 90^*°*^, while on the right surface, the distribution shifts to angles less than 90^*°*^. This mirror symmetry indicates that the terminal regions of the IDRs project radially outward from the condensate surfaces, maintaining an extended conformation that facilitates solvent exposure and potential interaction with surrounding molecules, consistent with a previous study.^82^

Under conditions of high repulsive crowding (Figure 3C), TDP-43 CTD condensates exhibit pronounced compaction. The spatial probability distributions of all three segments shift inward along the *z*-axis, reflecting a denser internal architecture and reduced overall condensate volume. Notably, the *α*-helical region becomes more strongly centralized, exhibiting a marked enrichment at the condensate core. Its orientation distribution also becomes narrower, suggesting a more ordered and preferential alignment relative to the *z*-axis. This behavior likely arises from enhanced Helix-Helix interactions, which are energetically favored under high-density conditions that promote molecular packing. However, the observed compaction is primarily driven by excluded-volume effects: repulsive crowders in the surrounding dilute phase exert steric pressure that effectively “squeezes” TDP-43 CTD chains into the dense phase. This entropic confinement favors configurations that reduce the interfacial area between protein-rich and crowder-rich phases, thereby minimizing the total free energy. As a result, while the IDRs remain more peripherally distributed compared to the helix, their spatial segregation becomes more pronounced than in the crowder-free case. Together, these findings underscore how purely entropic forces imposed by repulsive crowding govern condensate organization. The system adopts a compact, layered architecture in which structured regions like the *α*-helix are centralized for optimal packing, while disordered IDRs are compressed toward the surface. This arrangement enhances condensate stability by integrating tight core packing with retained peripheral flexibility, preserving dynamic properties at the condensate interface.

In contrast, the introduction of attractive crowders gives rise to a swollen, more sparse condensate structure (Figure 3D). The spatial distribution profiles reveal that both IDR1 and IDR2 now explore a wider range of positions throughout the condensate and frequently extend into the dilute phase, indicating the formation of an expanded and softened interfacial layer. The *α*-helix, by comparison, exhibits a preferential localization near the condensate surface, likely driven by favorable enthalpic interactions with crowders that selectively associate with this compact, high-contact-density region. Within the condensate, the orientation distribution of the *α*-helix becomes broader and more isotropic compared to repulsive or crowder-free conditions, though it still displays a tendency to align parallel to the interface. These observations suggest that attractive crowders compete with Helix-Helix contacts that previously stabilized the condensate interior. Rather than promoting compaction, crowders disrupt the native interaction network, effectively solvating surface residues, reducing internal packing density, and promoting condensate expansion. Notably, the *α*-helix becomes enriched at two opposing interfacial zones, indicating selective partitioning into regions where crowder interactions are strongest. Meanwhile, the IDRs adopt extended conformations that bridge spatially distant regions, reinforcing their role as flexible linkers connecting structured domains. This emergent architecture, which features stable interaction hubs formed by structured elements and dynamic connectors provided by disordered regions, mirrors a general organizing principle observed in diverse biomolecular condensates.^82^ Angular distribution analysis reveals that the *α*-helix maintains a preferentially parallel orientation at the interface even in the presence of attractive crowders, whereas both IDRs exhibit outward-facing termini indicative of a radially extended configuration. While the orientational bias of the *α*-helix becomes less pronounced under attractive crowding, likely due to enhanced spatial dispersion, the overall directional tendencies of all regions persist. This observation highlights the robustness of intrinsic region-specific conformational preferences, which remain largely preserved despite significant enthalpic perturbations introduced by attractive interactions.

### Region-specific structure-dynamics decoupling of TDP-43 CTD within condensates

To elucidate the region-specific interaction dynamics that govern phase separation, we quantified the kinetics of inter-chain contacts within TDP-43 CTD condensates. Prior studies have shown that condensates exhibit markedly slower relaxation dynamics compared to dilute phases.^69^ Building on this foundation, we developed a quantitative framework to dissect the microscopic interaction patterns among distinct TDP-43 CTD segments (Figure 4A; see Materials and Methods for details). Our analysis centers on two complementary metrics: (1) the inter-chain region-based contact number 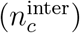, which reflects the thermodynamic likelihood of sustained contact between two segments, and (2) the contact relaxation time (*τ*), which captures the timescale over which those contacts persist. Specifically, 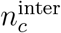 is defined as the number of inter-chain contacts per residue pair between two defined regions of TDP-43 CTD, while *τ* is extracted from the autocorrelation function of contact presence, with larger values indicating slower relaxation and more stable interactions. This dual-metric approach enables a systematic comparison of contact strengths and lifetimes between the structured *α*-helix and the flanking IDRs under varying crowding conditions.

**Figure 4:**
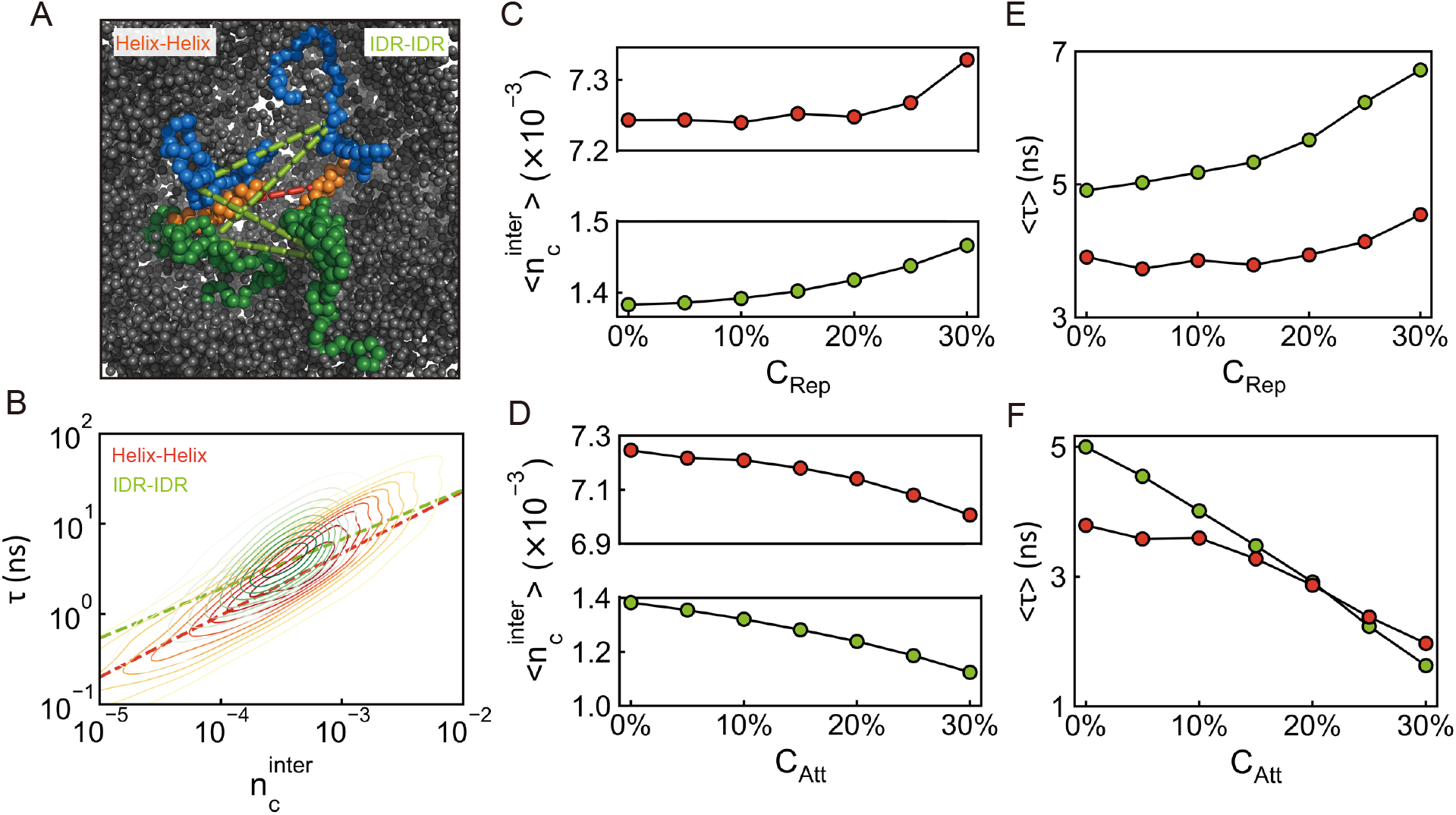
Region-specific structural and dynamical characteristics of TDP-43 CTD within condensates under different crowding conditions. (A) Schematic representation illustrating inter-chain region-based contacts involving structured (*α*-helix, red) and disordered (IDRs, blue and green) segments of TDP-43 CTD. (B) Two-dimensional contour plot showing the relationship between inter-chain region-based contact number 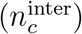 and contact relaxation time (*τ*), distinguishing regimes associated with Helix-Helix and IDR-IDR contacts. Dashed lines indicate linear fits for each interaction type. (C, D) Average inter-chain region-based contact number 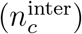 as a function of crowder concentration for (C) repulsive and (D) attractive crowders. (E, F) Mean contact relaxation time (*τ*) as a function of crowder concentration for (E) repulsive and (F) attractive crowders.

Our computational analysis of the two-dimensional 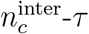 contour maps reveals key features of interdomain interactions within TDP-43 CTD condensates (Figure 4B). First, across all interaction types, we observed a strong positive correlation between inter-chain contact number 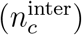 and contact relaxation time (*τ*), indicating that more frequent contacts are associated with slower, more persistent dynamics. Second, distinct interaction classes occupy well-separated regions in the 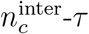 landscape: Helix-Helix cluster at high 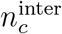 and low *τ*, reflecting strong interactions and relatively fast dissociations. By contrast, IDR-IDR contacts exhibit lower contact numbers and longer lifetimes, consistent with weak, persistent interactions. This systematic separation underscores the differential roles of structured and disordered domains in modulating condensate cohesion: while helices serve as stable interaction hubs but the interactions can be dynamically modulated, the IDRs contribute weak connectivity that facilitates molecular fluidity.

Our quantitative analysis of the mean inter-chain contact number 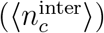 and mean contact relaxation time (⟨ *τ⟩*) reveals a systematic modulation of TDP-43 CTD interaction dynamics by crowding agents (Figures 4C-F, S10-S14). The positions of different interaction types within the 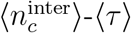 parameter space shift coherently in response to both repulsive and attractive crowders, demonstrating that the physicochemical properties of the surrounding environment directly tune the strength and lifetime of intermolecular interactions. These trends suggest a potential cellular mechanism for regulating the material properties of condensates by tailoring the crowding milieu to modulate inter-chain contact networks and dynamics.

In repulsive crowding environments, both 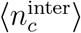 and ⟨*τ⟩* increase concurrently, indicating that inter-chain interactions become simultaneously stronger and more kinetically stable (Figure 4C). This behavior is consistent with the compaction observed in condensates under excluded-volume pressure, where tighter molecular packing reinforces contact persistence. In contrast, attractive crowders induce the opposite response: 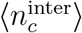 and ⟨*τ⟩* both decrease with crowder concentration, reflecting weaker, more transient inter-chain associations and a more diffuse condensate structure (Figure 4D). These opposing trends underscore how crowding agents, depending on their interaction specificity, can bidirectionally regulate both the thermodynamic and kinetic dimensions of condensate organization in a concentrationdependent manner.

Our region-specific analysis further uncovers striking differences in the interaction dynamics among TDP-43 CTD domains. Helix-Helix contacts, although thermodynamically the stronger, exhibit unexpectedly faster kinetics under repulsive and low attractive crowder concentrations, than IDR-IDR contacts (Figures 4E-F). In contrast, IDR-IDR contacts, despite their weaker binding affinity, display significantly longer relaxation times, suggesting the formation of persistent, long-lived interactions. This apparent paradox stems from the intrinsic structural properties of each segment: the conformational flexibility of IDRs allows them to dynamically reconfigure and maintain inter-chain interactions even under fluctuating conditions, whereas the rigidity of the *α*-helical region limits such adaptability, resulting in more transient contacts despite stronger binding forces. These differences underscore the specialized functional roles of structured versus disordered regions in maintaining condensate cohesion under dynamic conditions.

At high concentrations of attractive crowders, we observed a pronounced inversion in interaction behavior (Figure 4F). Helix-Helix contacts not only remain the strongest in terms of binding affinity but also become the most kinetically stable interactions. This shift likely arises from preferential stabilization of compact helical domains by attractive crowders, which reduce interfacial fluctuations and promote persistent contact configurations. Concurrently, the decreased condensate density enables IDRs to adopt extended conformations and form transient, rapidly exchanging inter-chain contacts. This complementary behavior with rigid helices providing stable structural cores and flexible IDRs facilitating dynamic connectivity, reveals a compensatory mechanism by which condensates balance architectural integrity and molecular fluidity. These findings highlight how the division of labor between structured and disordered domains equips condensates with the ability to dynamically adapt to diverse and changing physicochemical environments.

Our analysis reveals fundamentally distinct interaction and dynamic paradigms between structured and disordered regions within biomolecular condensates. Helix-Helix interactions are thermodynamically strong but kinetically transient, frequently breaking and reforming as the condensate undergoes internal reorganization. In contrast, IDR-IDR interactions, although weaker in binding affinity, exhibit remarkable persistence, sustaining long-lived contacts over extended periods. This dichotomy stems from the intrinsic conformational plasticity of IDRs, which allows them to dynamically adapt their configurations and maintain interactions even under fluctuating physicochemical conditions. ^83^ The complementary properties of these domains with rigid helices acting as stable anchoring nodes and flexible IDRs enabling dynamic inter-chain connectivity, may suggest a mechanistic rationale for the evolutionary conservation of disordered regions in phase-separated systems. The ability of IDRs to mediate long-range associations while permitting internal fluidity and rearrangement appears essential for fulfilling the dual requirements of condensate stability and dynamic adaptability, as the hallmarks of functional biomolecular condensates.^84^

## Discussion and Conclusions

Macromolecular crowding is increasingly recognized as a critical extrinsic factor capable of either promoting or inhibiting biomolecular phase separation, depending on the nature of interactions between the crowders and the phase-separating species. ^38,85,86^ In our simulations, repulsive (inert) crowders, interacting exclusively via excluded-volume effects, robustly promoted phase separation of the TDP-43 CTD. These crowders increased the protein density within condensates and enhanced protein partitioning into the dense phase. This result aligns with theoretical predictions of depletion-induced phase separation and is consistent with prior experimental observations; for instance, inert polymers such as PEG have been demonstrated to enhance protein condensation (e.g., FUS) and can even induce transitions from liquid-like to gel-like states at sufficiently high concentrations.^52,67^ Under repulsive crowding conditions, we found strong and predictive correlations among molecular-scale observables: single-chain compaction (*R*_*g*_), dimer contact number 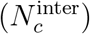, and condensate density (*ρ*_*h*_). This consistency allows inference of condensate formation propensity directly from individual molecular properties, indicating a coherent physical picture dominated by volume-exclusion-driven compaction (Figure 5). In contrast, attractive crowders, which directly interact with the TDP-43 CTD through soft-binding interactions, suppressed phase separation by solvating the protein chains and disrupting inter-protein contacts. Our simulations showed that attractive crowders still maintained strong predictive relationships among *R*_*g*_, 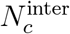, and *ρ*_*h*_ (Figure 5), though with quantitatively distinct scaling behaviors. These concentration-dependent modulations reflect competition between enthalpic protein-crowder interactions and entropic protein-protein interactions, leading to significant alterations in the energy landscape governing condensate formation.^55^ Similar outcomes have been reported in CG simulations of other LLPS systems, where even mildly attractive cosolutes disrupt phase behavior by masking or competing with homotypic protein-protein interactions. ^55,87^

**Figure 5:**
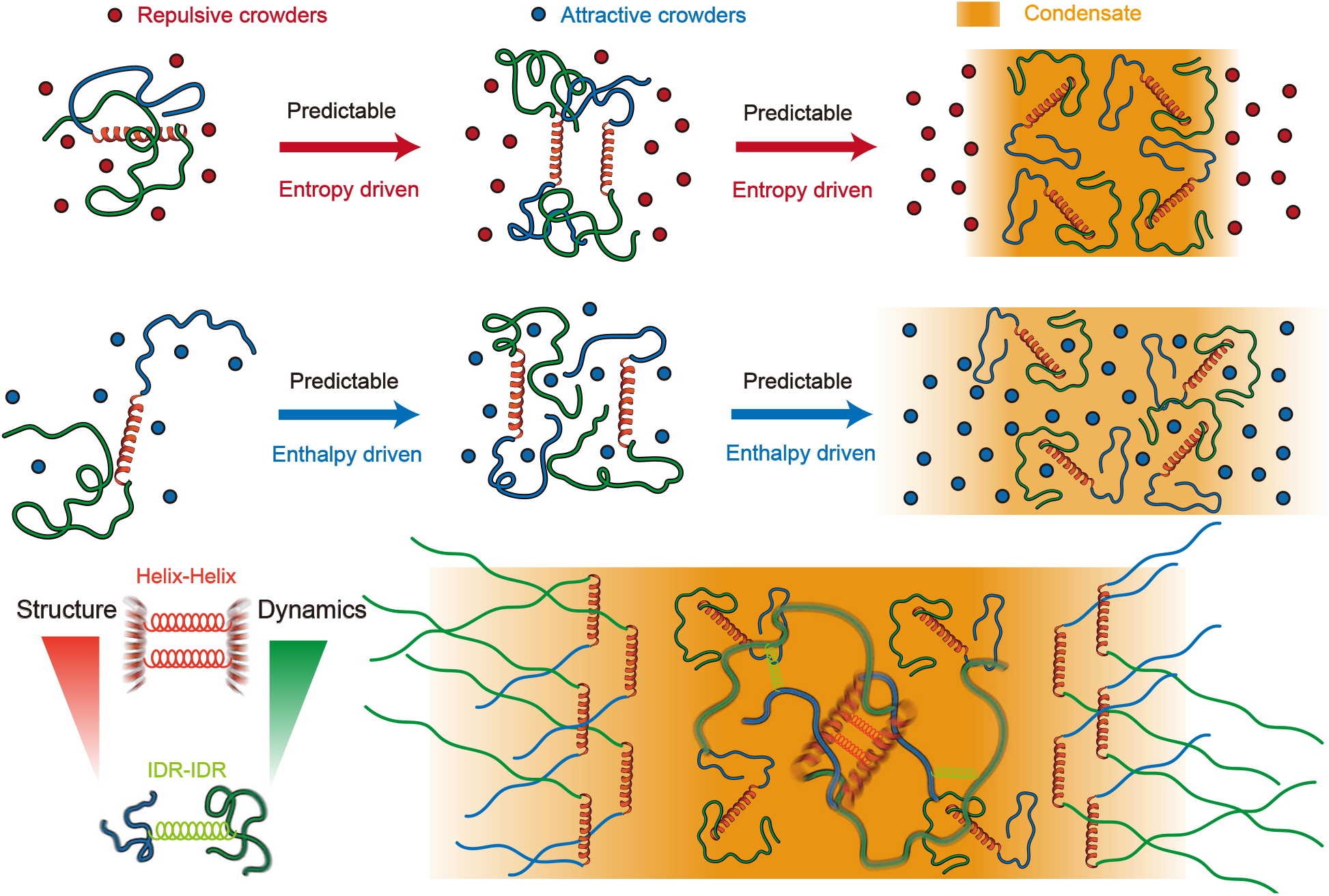
Schematic illustration summarizing the key findings of this study. Under both repulsive and attractive crowding conditions, strong correlations exist among single-chain compaction, inter-chain dimer contacts, and the resulting condensate density for TDP-43 CTD. Within phase-separated condensates, structured *α*-helical domains preferentially localize to the condensate interior, forming stable Helix-Helix interactions that enthalpically stabilize the condensate. These helices typically adopt orientations parallel to the condensate surface. In contrast, IDRs extend outward from the condensate surface, contributing entropically to interfacial flexibility. Notably, the inter-chain structural interaction strength and dynamic contact persistence exhibit region-specific correlation: helices form strong but dynamic interactions, while IDRs engage in weaker yet more persistent contacts. This division underpins both the structural stability and dynamic adaptability of TDP-43 CTD condensates.

A striking outcome of our simulations is the spatial segregation of the TDP-43 CTD’s *α*-helical segments from its disordered regions within condensates (Figure 3). We found that the TDP-43 CTD’s short helix-forming region tends to self-associate, clustering with helices from other molecules and forming a semi-ordered, dynamic network, whereas the flanking IDRs are less dense with slow relaxation. Such internal self-organization is conceptually akin to the scaffold-client paradigm described for multi-component biomolecular condensates. ^88,89^ In our case, the helical segments act analogously to scaffolds, providing highly interactive surfaces that concentrate together, while the surrounding disordered segments behave like client regions. The net result is a condensate with an internal helix-rich core with IDR-rich shell architecture, rather than a perfectly homogeneous droplet. Indeed, Conicella et al. showed that two glycine residues in this helix normally limit its extension, and mutating them to more helix-promoting amino acids dramatically enhanced TDP-43 phase separation and reduced its droplet fluidity. ^25^ Those results underscore that TDP-43 CTD helix is a tunable module for self-association, consistent with our simulations: when the helix is allowed to stabilize and cluster modulated by increasing the number of repulsive crowders, TDP-43 molecules condense more readily and organize into a less uniform, more biphasic structure. Thus, our work provides a molecular-scale picture for how region-specific interactions can give rise to internal condensate architecture, which is a physical principle that may generalize to other multi-domain proteins and help explain the formation of layered or core-shell structures observed in cellular condensates.^90–92^

Our analysis of intra-condensate dynamics reveals a pronounced asymmetry in the interaction kinetics of structured and disordered regions within TDP-43 CTD condensates, implications for their material state. Specifically, Helix-Helix contacts are short-lived but thermodynamically strong, while IDR-IDR interactions are weaker yet long-lived and dynamically persistent (Figure 5). This duality gives rise to a heterogeneous dynamic landscape: the *α*-helical regions form a fast-relaxing, percolated scaffold that imparts mechanical rigidity, whereas the IDR-rich regions preserve high mobility and fluidity. Such behavior is emblematic of a “network fluid” or viscoelastic gel, which is a phase-separated system that exhibits both solid-like and liquid-like characteristics depending on the timescale of observation.^93–95^ This emergent biphasic behavior is consistent with recent theoretical and experimental insights into condensate rheology that the viscoelasticity of biomolecular condensates arises from the balance between transient, multivalent interactions and polymeric flexibility.^84^ Furthermore, it has been demonstrated that condensate dynamics are governed by two principal factors: (1) specific, often sequence-encoded inter-chain interactions, and (2) entropic contributions from polymer length and conformational freedom.^96^ In our model, the helix provides factor (1) through strong, compact contacts, while the flexible IDRs fulfill factor (2) by supporting extensive conformational sampling and adaptive contact networks. The interplay of these features enables TDP-43 CTD condensates to span a rheological continuum from fluidlike to gel-like states. Our results also offer a mechanistic perspective on how condensate material properties can be tuned. For example, enhancing the stability of the helical domain, either through helix-promoting mutations or post-translational modifications,^97^ would be expected to increase the persistence of inter-helical contacts, shifting the condensate toward a more solid-like state. Conversely, disruption of Helix-Helix interactions, or promotion of IDR solvation via binding partners or osmolytes, could fluidize the condensate by reducing network connectivity. These predictions align with experimental findings in TDP-43 and other systems, where condensate mobility is reduced by strengthening interaction motifs or by applying crowding conditions that promote network formation.^25,67,98^ Altogether, our findings highlight how microscopic kinetic asymmetries between domains underlie emergent macroscopic material states, a principle likely generalizable to other modular, multidomain proteins involved in phase separation.

We acknowledge that our simulations focused exclusively on the CTD of TDP-43, omitting contributions from the full-length protein, including its NTD and RRMs, and interactions with RNA or other cellular partners, elements known to modulate phase behavior significantly.^99^ Additionally, the macromolecular crowders in our model were implemented as generic, static particles, serving as a simplified representation of the crowded cellular milieu.^48,100^ In reality, cellular crowding arises from a heterogeneous mixture of macromolecules, including inert polymers, multivalent phase-separating proteins, and other active components that can partition into condensates or modulate their material properties through specific interactions. ^101^ Furthermore, the CG models employed here, while enabling large-scale exploration of phase behavior, inherently lack the resolution and temporal reach to capture long-timescale phenomena such as condensate aging, maturation, or pathological fibrillization that frequently occur *in vivo*, especially under stress conditions.^102^ Future developments could incorporate these additional layers of biological complexity to refine our understanding of TDP-43 phase separation. For example, introducing ATP-dependent chaperones, posttranslational modifications, or ligand-induced conformational switching into the simulation framework may reveal how cells dynamically regulate condensate composition, turnover, and pathological progression. ^37,103^ Ultimately, a comprehensive model of TDP-43 behavior will require integration of structural domains, multicomponent biochemical interactions, and active cellular regulation, highlighting the importance of bridging minimal physical models with biologically realistic contexts.

In summary, this study presents a theoretically grounded and integrative framework for elucidating biomolecular phase separation in crowded, cellular-like environments. By systematically contrasting repulsive and attractive crowding regimes and disentangling the distinct roles of the structured *α*-helix and disordered regions (IDRs) within the TDP-43 CTD, we uncover key physical principles that govern condensate formation, internal architecture, and dynamic behavior. Our results show that repulsive crowders promote compaction and robust phase separation through entropic depletion forces, whereas attractive crowders reshape condensate morphology and dynamics via enthalpic interactions that can override intrinsic molecular interaction hierarchies. Crucially, we reveal a division of labor between domain types: Helix-Helix contacts are strong yet transient, enabling structural anchoring, while IDR-IDR contacts are weaker but long-lived, ensuring connectivity and dynamic flexibility. This synergy between structure and disorder bridges molecular-scale interactions and mesoscale condensate properties, shedding light on how cells modulate the composition, organization, and material state of membraneless organelles through physicochemical cues.^84,88^ Although focused on TDP-43, the principles uncovered here have broad applicability to diverse phase-separating proteins and offer a conceptual basis for understanding the assembly, regulation, and pathological misregulation of biomolecular condensates across biological systems.

## Data availability

The necessary files for setting up LAMMPS simulations and analysis programs/scripts are publicly available at GitHub with link: https://github.com/GuoqingZhang1/LLPScrowder.git. This repository includes the following components:

- Simulation Files: LAMMPS input files for running the molecular simulations.
- Analysis Tools: Python programs/scripts for analyzing simulation data.

## Acknowledgments

This work was supported by the National Natural Science Foundation of China (Grant Nos. 12474201 and 32201020), the General Program of the Guangdong Basic and Applied Basic Research Foundation (Grant No. 2024A1515010862), and the Guangdong Provincial Project (Grant No. 2023QN10×037). The authors also acknowledge the Green e Materials Laboratory (GeM) and HPC+AI Intelligence Computing Center at the Hong Kong University of Science and Technology (Guangzhou) for providing computational support.

## Supporting Information Available

- Supporting Information: figures S1-14 for additional results.

This material is available free of charge via the Internet at http://pubs.acs.org/.

## Graphical TOC Entry

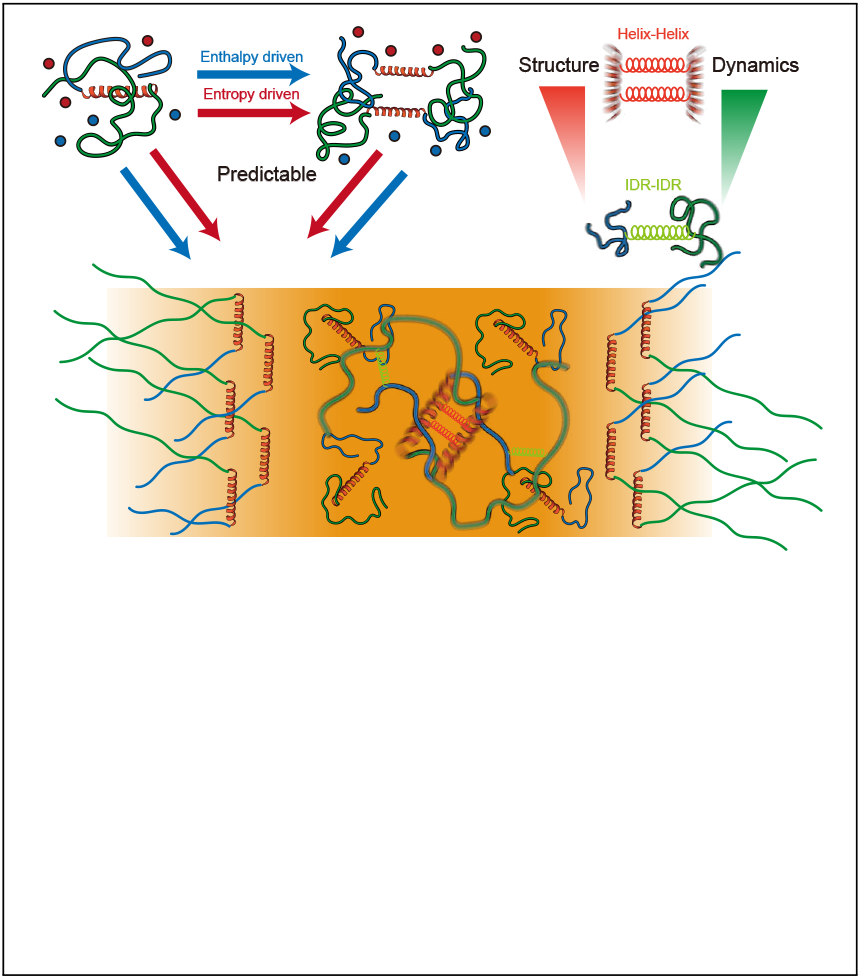

