## Supplementary material for "Repulsive vs. Attractive Crowding Distinctly Regulate TDP-43 Condensates through Region-Specific Structural Dynamics": SI Figures

for

### Additional figures

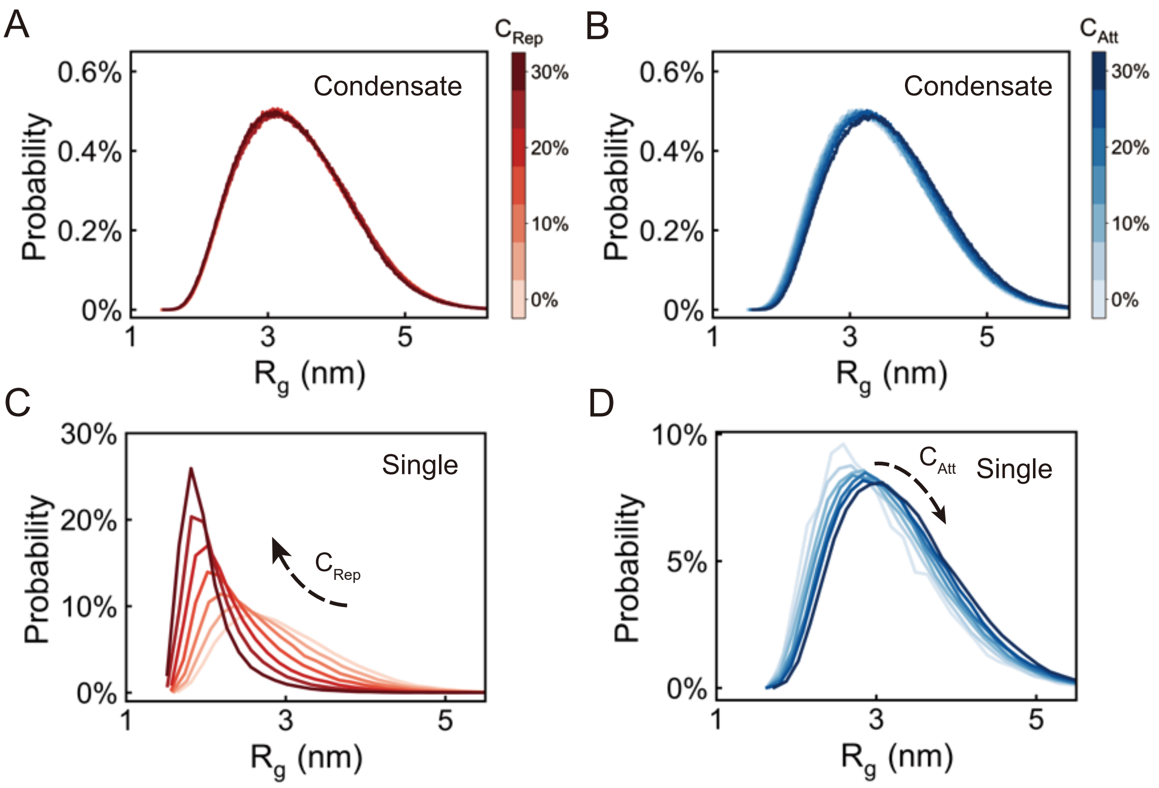


Figure S1: Distributions of the radius of gyration ($R_g$) of TDP-43 CTD under different crowding conditions. (A, B) $R_g$ distributions for TDP-43 CTD chains within condensates in the presence of (A) repulsive and (B) attractive crowders under varying concentrations. (C, D) $R_g$ distributions for single-chain TDP-43 CTD in the presence of (C) repulsive and (D) attractive crowders under varying concentrations. Each curve represents the probability distribution of $R_g$ sampled from simulations at the specified crowder concentration.

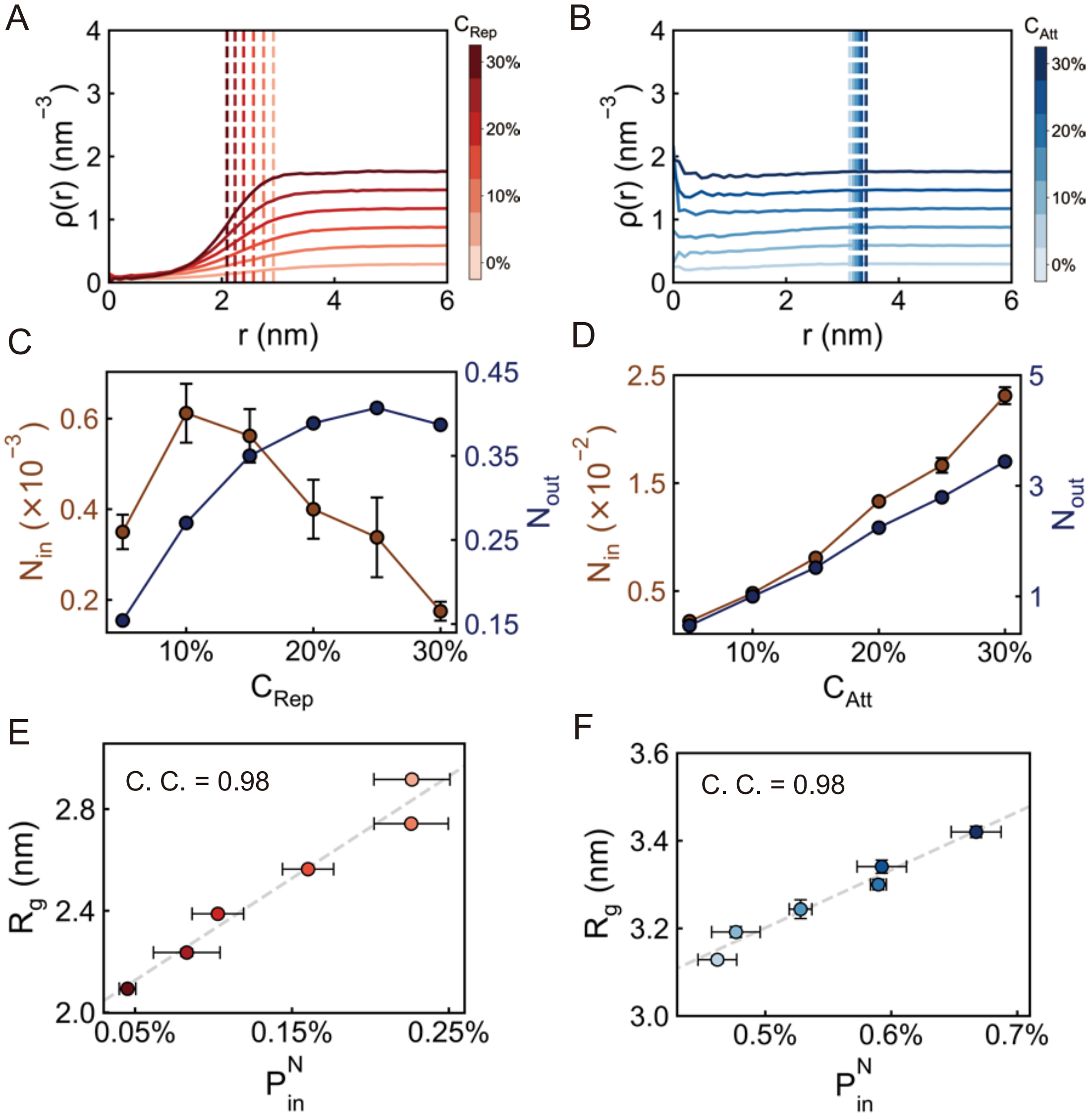


Figure S2: Radial density function (RDF) of crowders relative to the TDP-43 CTD center of mass in the single-chain state under different crowding conditions. (A, B) RDF profiles of crowders in the presence of (A) repulsive and (B) attractive crowders under varying concentrations. Dashed lines denote the average $R_g$ regions of the TDP-43 CTD chains under corresponding conditions. (C, D) Total number of crowders located inside ($N_{\mathrm{in}}$) and outside ($N_{\mathrm{out}}$) the average $R_g$ region of the TDP-43 CTD chain in the presence of (C) repulsive and (D) attractive crowders under varying concentrations. (E, F) Correlation between percentage of crowders located outside ($P_{\mathrm{in}}$) the average $R_g$ region of the TDP-43 CTD chain and the $R_g$ of the single TDP-43 CTD chain in the presence of (E) repulsive and (F) attractive crowders under varying concentrations. $P_{\mathrm{out}}$ is calculated as $N_{\mathrm{out}}/(N_{\mathrm{in}}+N_{\mathrm{out}})$ from (C, D). Dashed lines are provided as visual guides to highlight the trends with the correlation coefficients calculated. In panels (C-F), data points represent means $\pm$ standard errors computed from 5 equal partitions of the total dataset.

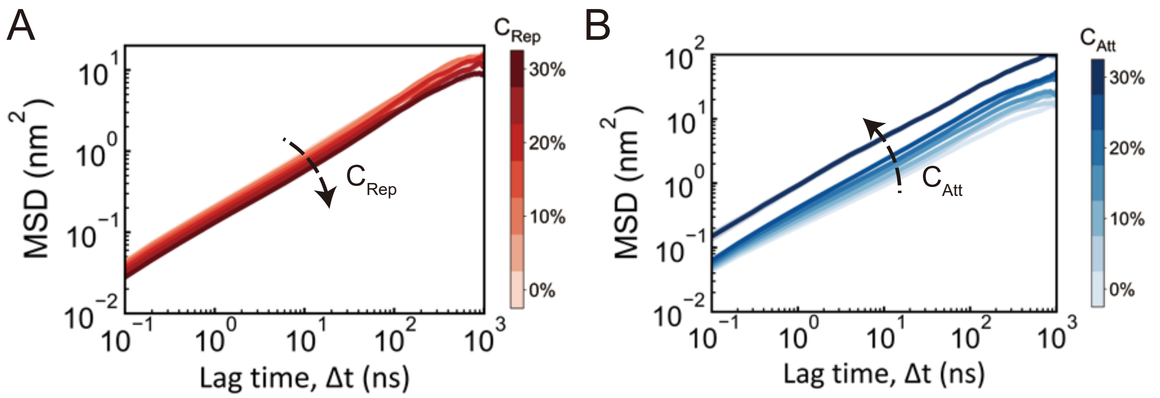


Figure S3: Mean square displacement (MSD) of TDP-43 CTD within condensates under varying concentrations of (A) repulsive and (B) attractive crowders. Data represent means ± standard errors computed from 5 equal partitions of the total dataset.

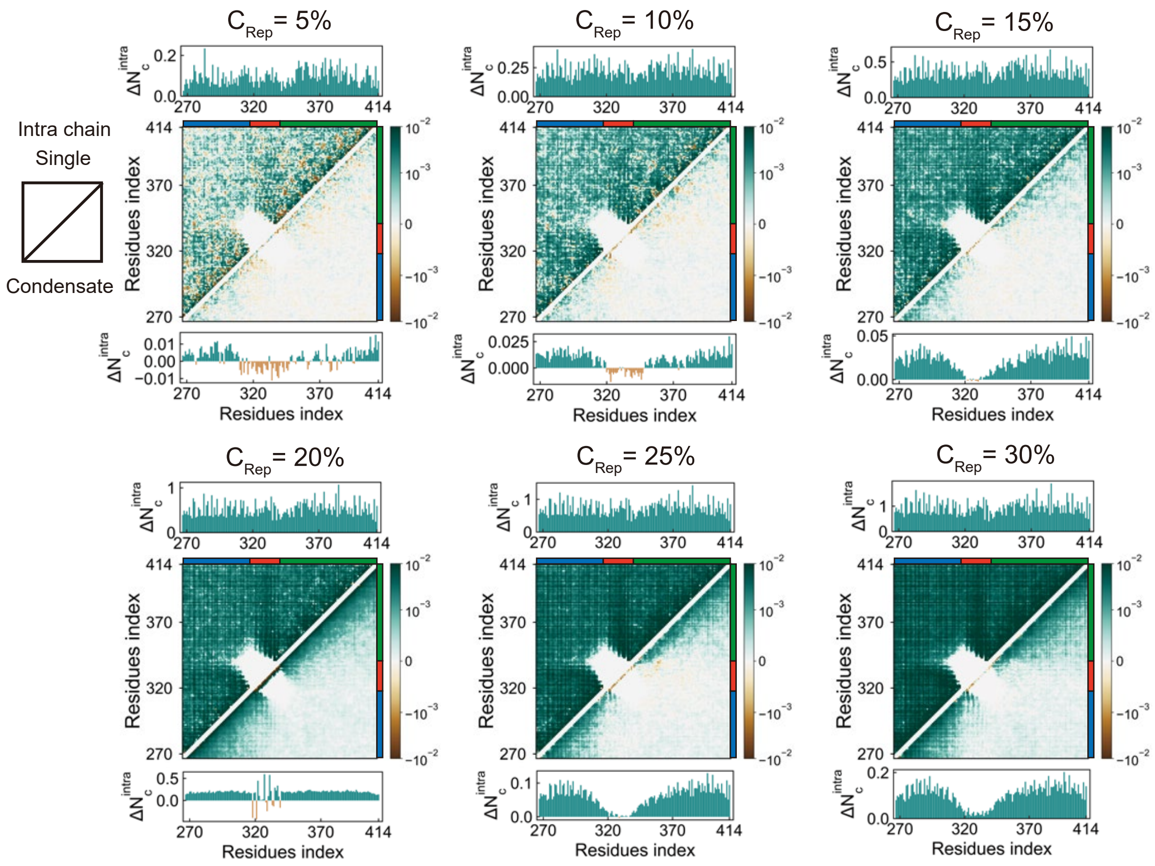


Figure S4: Differential intra-chain residue-residue contact probability maps of TDP-43 CTD under varying concentrations of repulsive crowders. Each map shows changes in contact probability relative to the crowder-free condition.

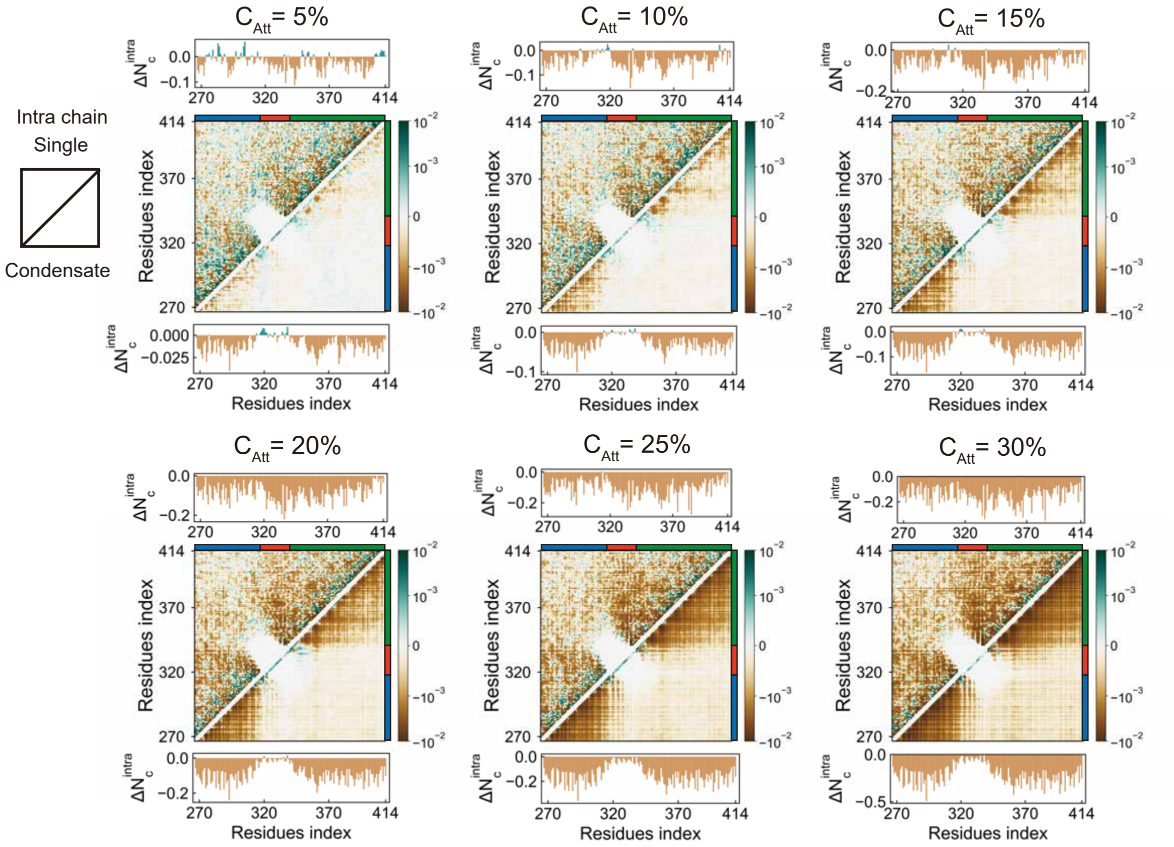


Figure S5: Differential intra-chain residue-residue contact probability maps of TDP-43 CTD under varying concentrations of attractive crowders. Each map shows changes in contact probabilities relative to the crowder-free condition, following the same format as Figure S4.

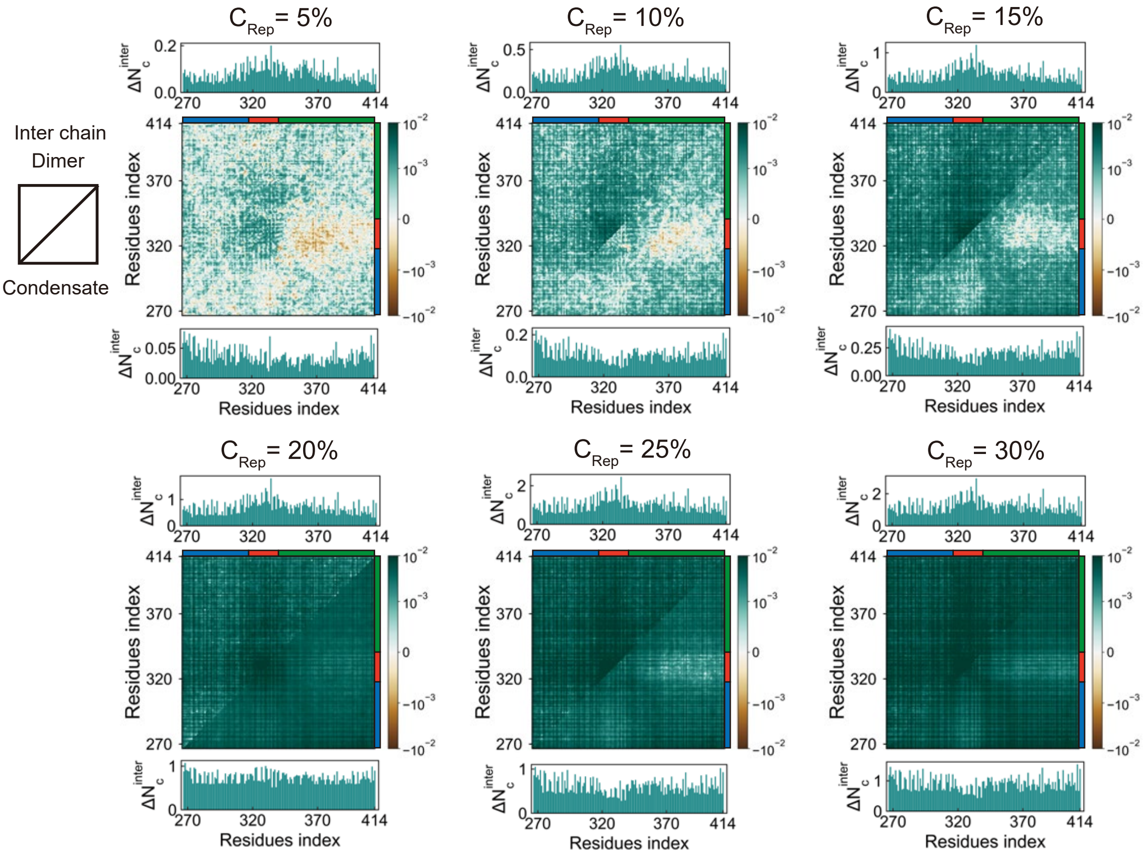


Figure S6: Differential inter-chain residue-residue contact probability maps of TDP-43 CTD under varying concentrations of repulsive crowders. Each map shows changes in contact probabilities relative to the crowder-free condition, following the same format as Figure S4.

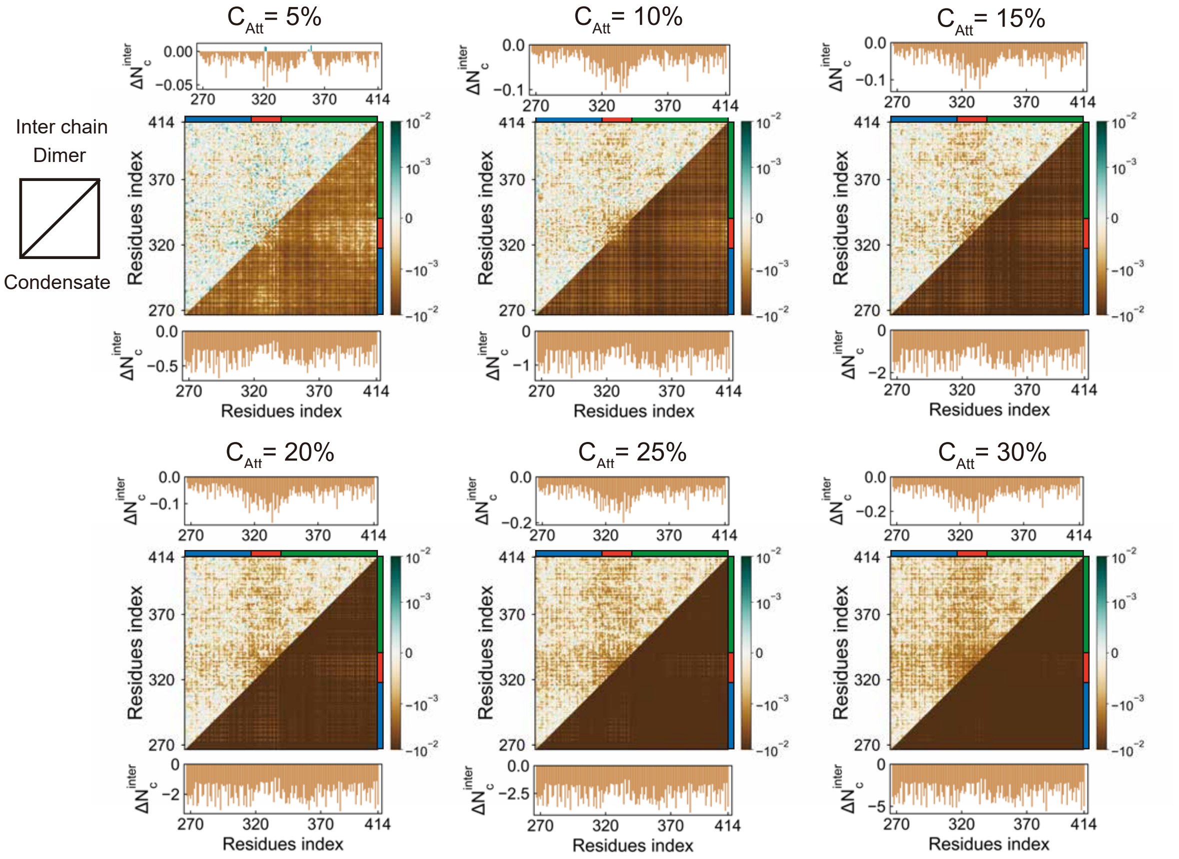


Figure S7: Differential inter-chain residue-residue contact probability maps of TDP-43 CTD under varying concentrations of attractive crowders. Each map shows changes in contact probabilities relative to the crowder-free condition, following the same format as Figure S4.

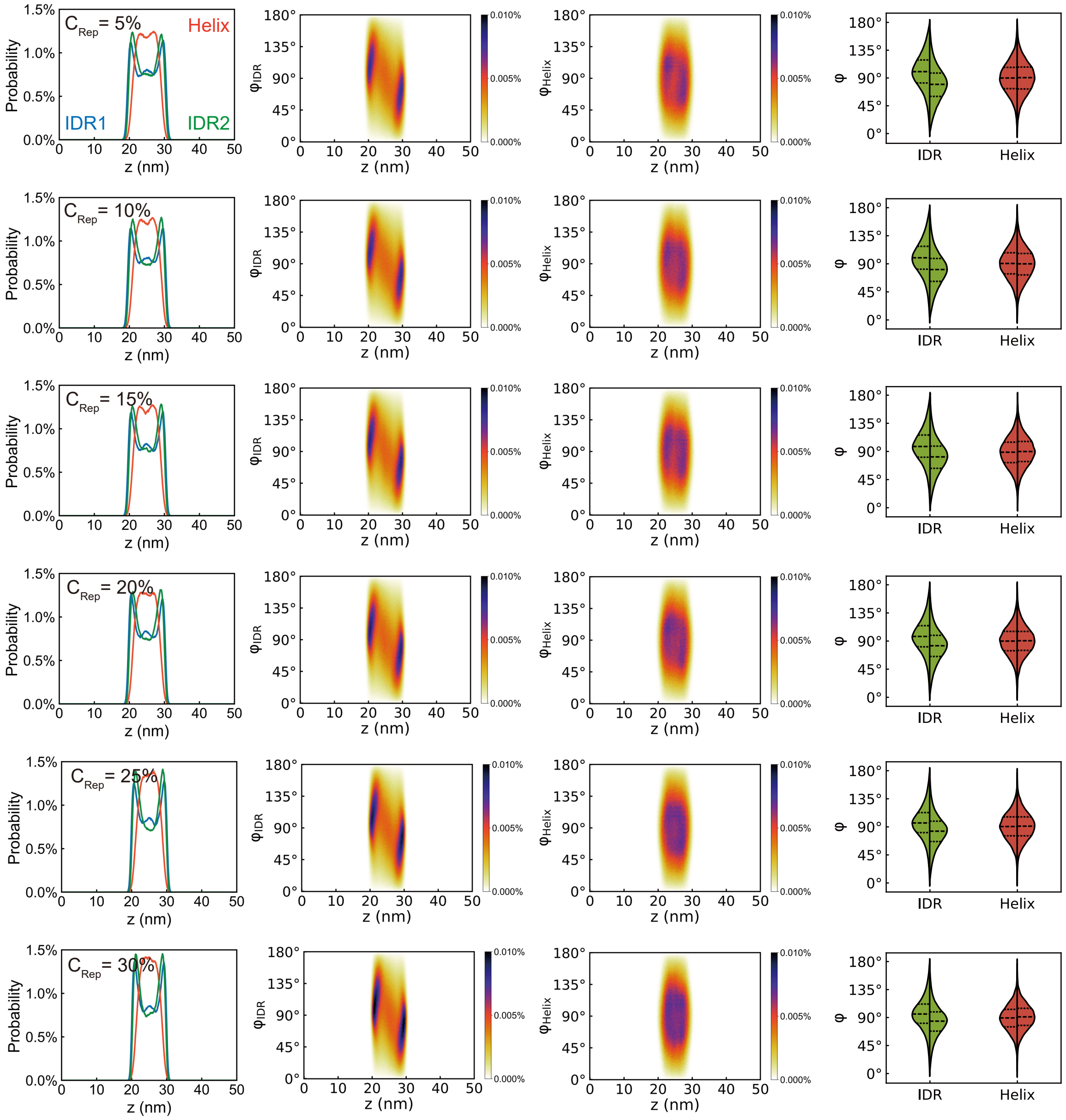


Figure S8: Region-specific spatial and orientational distributions of TDP-43 CTD segments within condensates under varying concentrations of repulsive crowders. Format and analysis are consistent with Figure 3 in the main text.

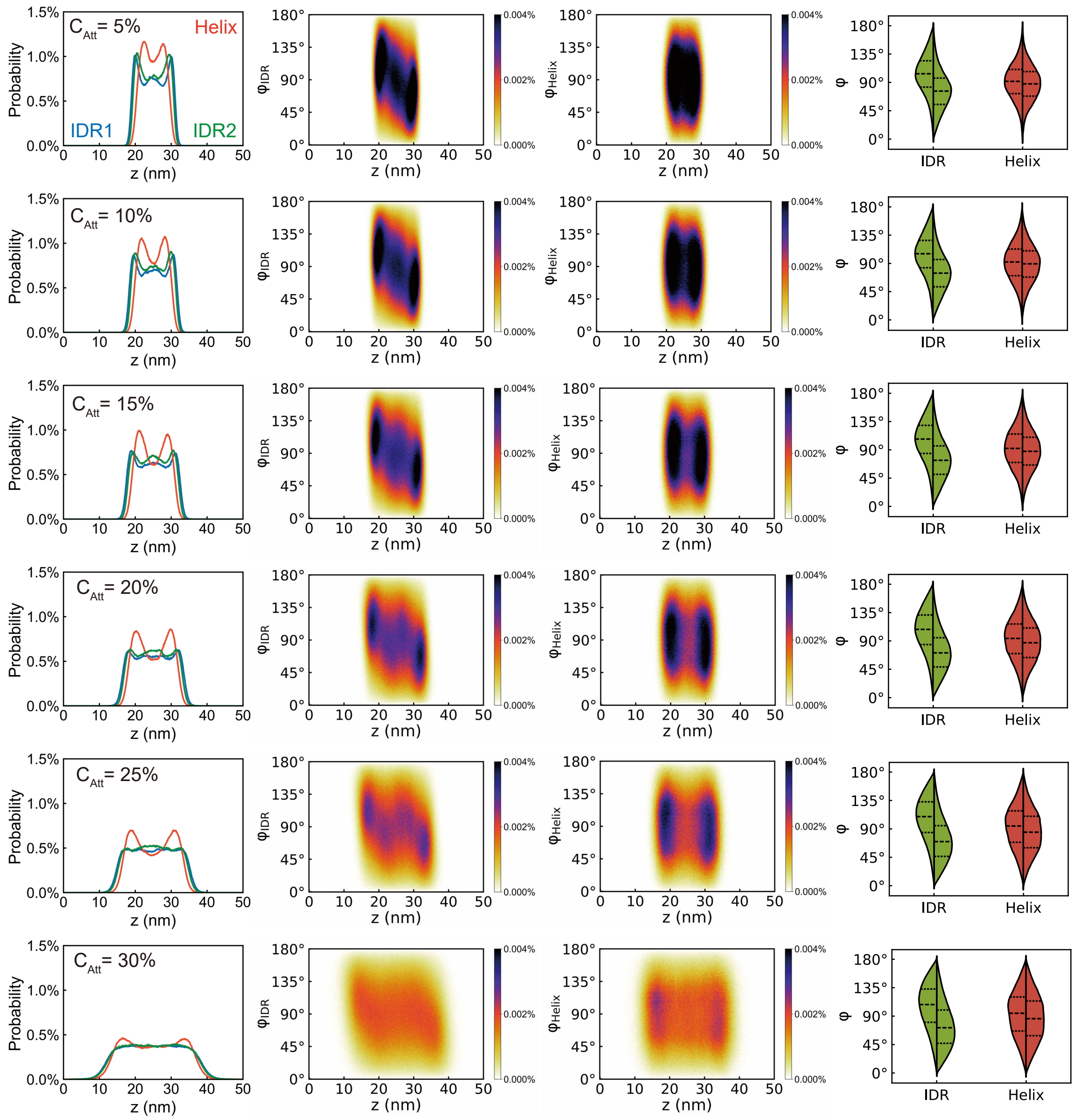


Figure S9: Region-specific spatial and orientational distributions of TDP-43 CTD segments within condensates under varying concentrations of attractive crowders. Format and analysis are consistent with Figure 3 in the main text.

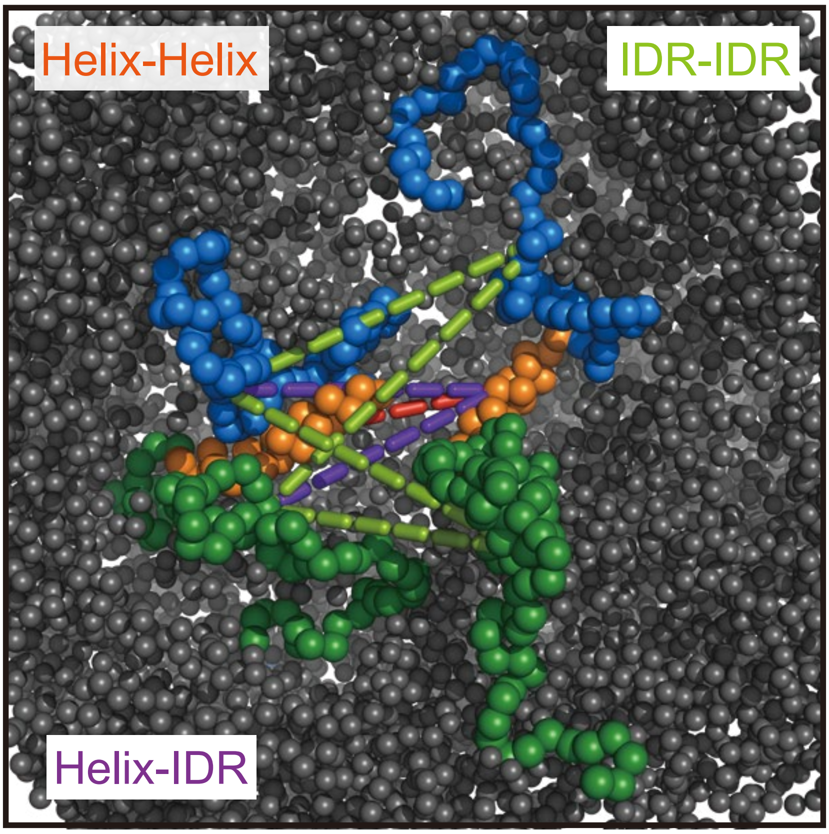


Figure S10: Schematic illustration of the three types of inter-chain region-based interactions analyzed in this study: Helix-Helix (orange), IDR-IDR (green), and Helix-IDR (purple). These interaction modes correspond to different combinations of the structured $\alpha$-helical region and the flanking IDRs of TDP-43 CTD.

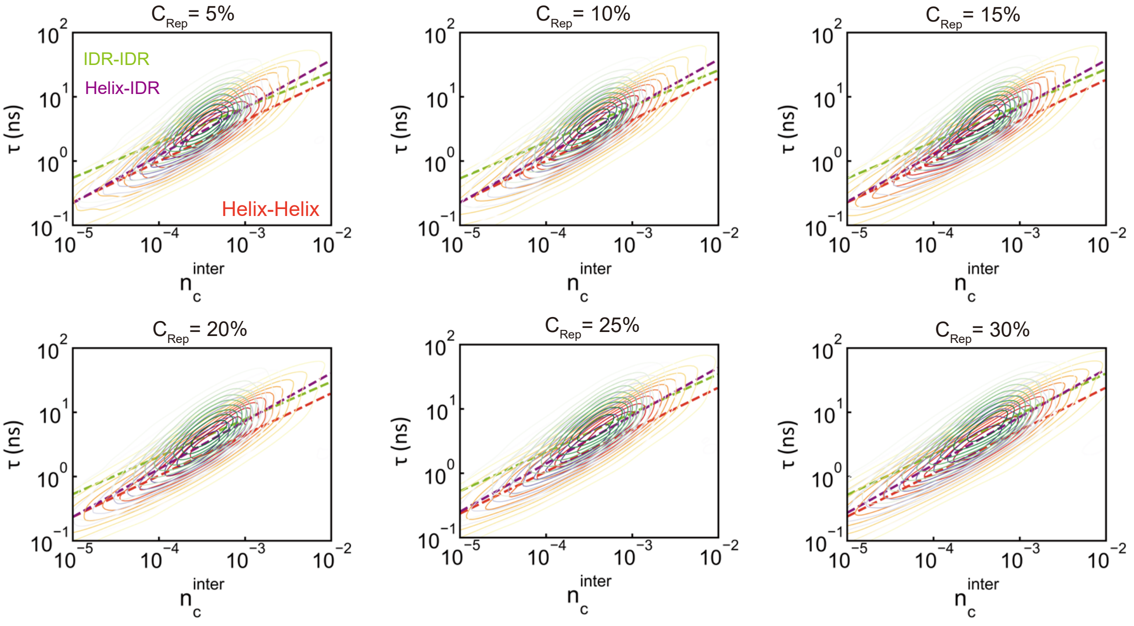


Figure S11: Two-dimensional contour plot showing the relationship between inter-chain region-based contact number ($n_c^{\text{inter}}$) and contact relaxation time ($\tau$) in the presence of repulsive crowders under varying concentrations. Dashed lines indicate linear fits for each interaction type. This analysis is equivalent to that shown in Figure 4B of the main text.

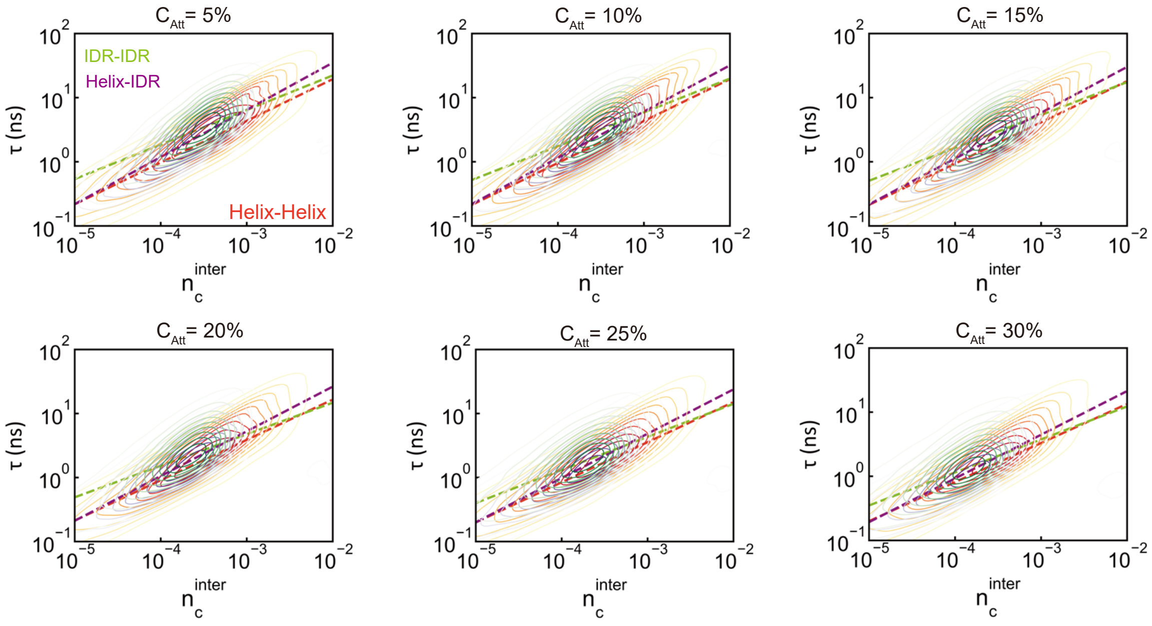


Figure S12: Two-dimensional contour plot showing the relationship between inter-chain region-based contact number ($n_c^{\text{inter}}$) and contact relaxation time ($\tau$) in the presence of attractive crowders under varying concentrations. Dashed lines indicate linear fits for each interaction type. This analysis is equivalent to that shown in Figure 4B of the main text.

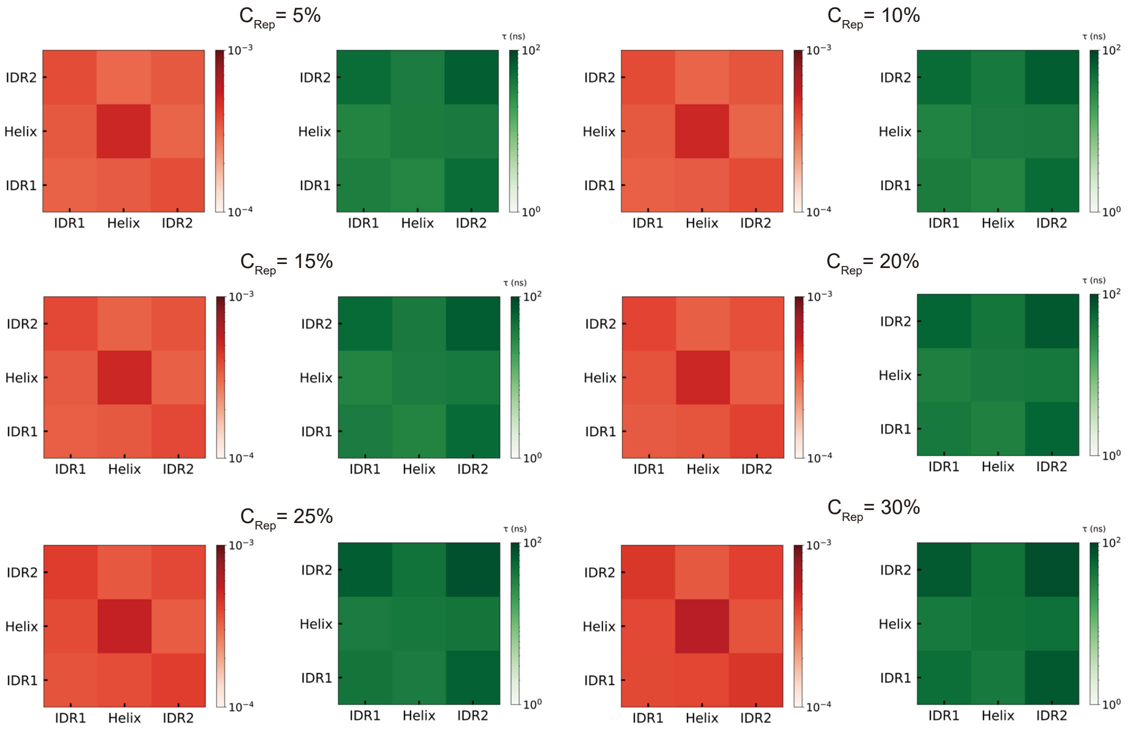


Figure S13: Heat maps showing the inter-chain region-based contact number ($n_c^{\text{inter}}$) and contact relaxation time ($\tau$) under varying concentrations of repulsive crowders.

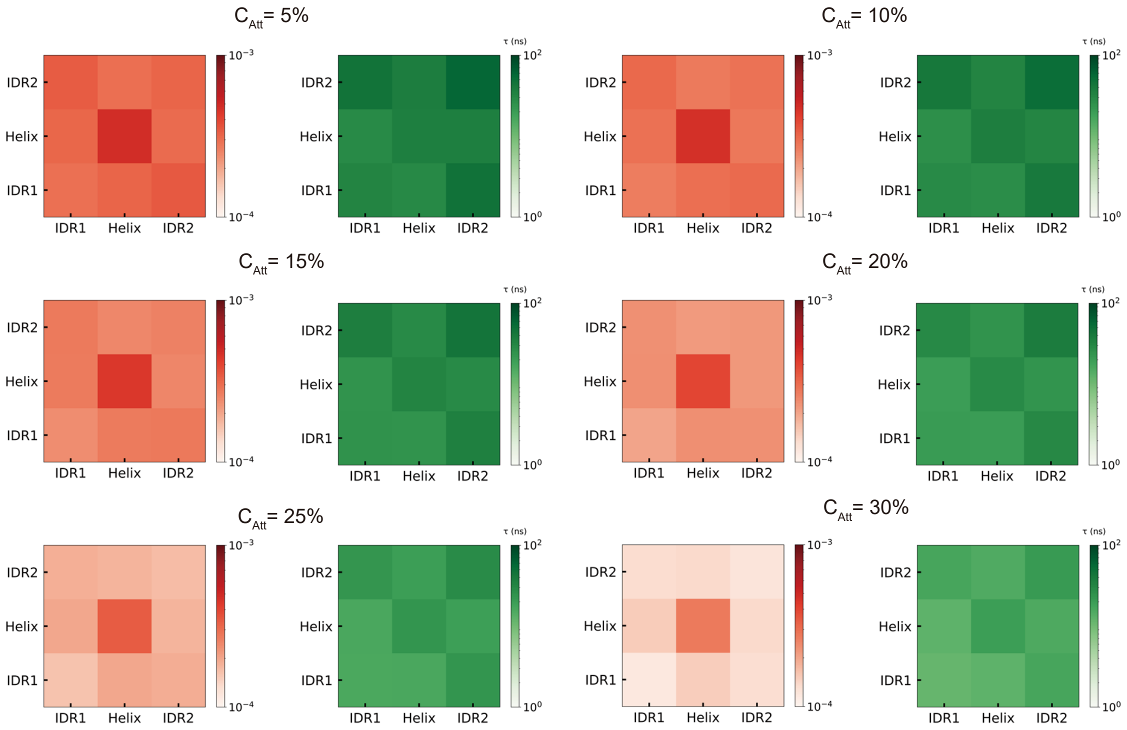


Figure S14: Heat maps showing the inter-chain region-based contact number ($n_c^{\mathrm{inter}}$) and contact relaxation time ($\tau$) under varying concentrations of attractive crowders.
